# Unlocking the Mycorrhizal Nitrogen Pathway Puzzle: Metabolic Modelling and multi-omics unveil Pyrimidines’ Role in Maize Nutrition via Arbuscular Mycorrhizal Fungi Amidst Nitrogen Scarcity

**DOI:** 10.1101/2023.10.13.562190

**Authors:** Bérengère Decouard, Niaz Bahar Chowdhury, Aurélien Saou, Martine Rigault, Isabelle Quilleré, Thomas Sapir, Anne Marmagne, Christine Paysant le Roux, Alexandra Launay-Avon, Florence Guerard, Caroline Mauve, Bertrand Gakière, Céline Lévy-Leduc, Pierre Barbillon, Pierre-Emmanuel Courty, Daniel Wipf, Bertrand Hirel, Rajib Saha, Alia Dellagi

## Abstract

Maize is currently the most productive cereal crop in the world (www.faostat.org). Maize can form a symbiotic relationship with the Arbuscular Mycorrhizal Fungus (AMF), *Rhizophagus irregularis*. In this relationship, the fungus provides the plant with additional water and mineral nutrients, while the plant supplies carbon compounds to the fungus. Little is known about the N metabolism disruption during symbiosis in both partners. To address this issue, two genetically distant maize lines were studied in terms of physiological and molecular responses to AMF inoculation by dual RNA-seq, metabolomics and phenotyping. Interestingly, the beneficial effects of the AMF were observed mainly under conditions of limited N fertilization. Under such conditions, the AMF helped maintain plant biomass production. The availability of nitrogen was found to be a crucial factor influencing all the traits studied showing that the level of N supply plays a pivotal role in determining how maize plants interact with the AMF. Despite the two maize lines showing different transcriptomic and metabolomic responses to *R. irregularis*, their agro-physiological traits remained similar. Both the plant and fungal transcriptomes were more significantly influenced by the level of N nutrition rather than the specific maize genotype. This suggests that N availability has a more profound impact on gene expression in both organisms than the genetic makeup of the maize plant. To understand the metabolic implications of this symbiotic relationship, we integrated transcriptomic data into our recently built multi-organ Genome-scale metabolic model (GSM) called iZMA6517. Remarkably, this modelling approach was supported by metabolomics profiling, in particular increased leaf pyrimidine levels in response to AMF inoculation under limiting N supply. Consistently, fungal genes involved in pyrimidine de novo synthesis and salvage were found to be expressed in symbiotic roots. Our work highlights nucleotide and ureides metabolism as previously unrecognized factors contributing to the symbiotic N nutrition facilitated by *R. irregularis*, thereby enhancing maize growth. This study demonstrates the effectiveness of integrating multi-omics approaches with mathematical modelling to uncover novel metabolic mechanisms associated with AM symbiosis, without a priori.

## Introduction

Nitrogen (N) is the fourth most abundant component of plant biomass, which explains its pivotal role in agriculture ^1,2^. Indeed, large inputs of N fertilizers are necessary to ensure optimal crop productivity ^1,3^. However, only 30 to 50% of the N available in the soil is taken up by most crops, resulting in important N losses which can cause damage to the environment such as water pollution and increased greenhouse effect ^4,5^. Ensuring crop productivity while minimizing N supply can be monitored by measuring nitrogen use efficiency (NUE). A better understanding of how to enhance NUE is a major challenge in the context of agroecology as it represents the capacity of the plant to produce grains or vegetative biomass per unit of available N ^3,6,7^. Direct uptake of minerals including N and P from the soil by plants is ensured by transporters known to drive the so called “direct transport”. Interestingly, to fulfil their N-associated metabolic needs, plants can establish symbiosis with beneficial soil microbes which help to improve their N nutrition by different mechanisms, some of which are well characterized. Exploiting the capacity of a crop to benefit from beneficial microbes in terms of N nutrition in order to maintain yields while reducing chemically synthesized N input is challenging in the context of a more sustainable agriculture ^8–13^. Nitrogen can be transferred from the soil to the roots when the plant is colonized by Arbuscular Mycorrhizal Fungi (AMFs), a symbiotic process occurring in 80% of land plants including cereals ^8,12–14^. The early steps of AMF symbiosis involve signal exchange between the host roots of the host plant and the fungus. This signalling process leads to establishment of a physical contact between the partners during which the fungus develops structures like a hyphopodium on the root epidermal cell followed by its growth inside these cells up to the cortical cells where it differentiates into highly branched structures called “arbuscules”. Around these arbuscules, the cortical cells undergo strong reorganization leading to the formation of a structure called “periarbuscular membrane” (PAM). Dual nutritional exchange occurs at the interface of this membrane where the transfer of carbon containing molecules derived from photosynthesis such as hexoses and lipids feed the fungus, which in turn provides minerals such as N and P ^8,11,15–17^. The extra-radical mycelium, which represents an extension of the arbuscular structures, allows the fungus to explore the surrounding rhizosphere and to take up minerals further translocated to the periarbuscular interface to be finally delivered to the host plant.

It is well established that phosphorus (P) deficiency is the main driver for the establishment of mycorrhizal symbiotic associations ^18–22^. In contrast, the benefit of a mycorrhizal symbiosis in terms of N uptake and utilization by the host plant remains to be fully characterized ^11–14^

Maize is currently the most productive cereal crop in the world (www.faostat.org) with a global yield exceeding that of rice ^23^. Different maize genotypes are able to develop a symbiotic association with AMF ^24,25^. Several studies have shown that AMF can supply substantial amount of N to maize ^26,27^. However, physiological and molecular characterization of maize’s interaction with AMF under different N supply levels has never been undertaken.

A Genome-Scale Model (GSM) encompasses the majority of known metabolic reactions within a biological system. It employs techniques such as flux balance analysis (FBA) ^28^, flux variability analysis (FVA) ^29^, and parsimonious FBA ^30^, to predict the flux of reactions accurately. Genome-scale metabolic modelling (GSM) is a computational tool to model plant metabolism and applied to plant system can highlight potential key metabolic pathways in plants in response to environmental cues ^31^. Initial efforts on maize metabolic models were limited to its leaf ^32^ and root ^33^ GSMs, which showed agreement with existing literature regarding phenotypical predictions. Most recently, a multi-organ GSM of maize line B73, iZMA6517 ^34^, is proposed which includes full scale GSMs of root, stalk, kernel, and leaf; connected via phloem vascular tissue. iZMA6517 is the largest maize multi-organ GSM of maize till date and captured accurate plant phenotype under heat and cold stress conditions.

In the present study, we have investigated in an integrated manner the impact of N nutrition on the interaction between two maize lines and the AMF *Rhizophagus irregularis* both at the physiological and molecular levels. Using phenotyping, dual RNA-seq transcriptomics and metabolomics approaches, we generated a set of paired data (same plants used for all analysis). We have characterized such interactions using two maize lines. Both maize lines displayed improvement of agronomic performance in response to inoculation with *R. irregularis* under limited N supply. Thereby, apart from experimental works, we used *i*ZMA6517 to predict plant agronomic performance and organ-specific metabolic bottlenecks accurately to explore the symbiotic relationship with *R. irregularis*. *i*ZMA6517 GSM identified pyrimidine biosynthetic pathway as a key physiological process involved in the efficiency of N utilization in the symbiotic association when N is limiting. Interestingly, this prediction was supported by monitoring pyrimidine levels in leaves of symbiotic plants. Thus, modelling approach highlighted nucleotide and ureides metabolism as previously unrecognized factors contributing to the symbiotic N nutrition facilitated by *R. irregularis*, thereby enhancing maize growth. This research highlights how the combination of multi-omics strategies and mathematical modelling can reveal new metabolic processes associated with AM symbiosis without preconceived notions. Our work highlights nucleotide and ureides metabolism as previously unrecognized factors contributing to the symbiotic N nutrition facilitated by *R. irregularis*, thereby enhancing maize growth.

## Material and methods

### Plant material, growth conditions and inoculation

The maize inbred lines used in this study B73 and Lo3 (Supplementary table 1). The reference maize genotype B73, for which the genome was fully sequenced, ^35^. The fungal strain used is the highest quality of purity strain *Rhizophagus irregularis* DAOM197198 (Agronutrition, Toulouse, France). Inoculum was prepared in distilled water and the spore suspension was added next to the germinated seed or plantlet with a pipette.

In all experiments, seeds were surface-sterilized as follows. Seeds were first incubated in ethanol for 5 minutes at 28°C followed by rinsing with distilled water. Then they were incubated for 45 minutes in a 15% commercial bleach solution with 0.01% Triton x100 then rinsed in distilled water. They were placed on wet sterile Whatman paper in Petri dishes closed with parafilm and incubated in the dark for 48-72h at 20°C. Germinated seeds were selected and then sown in pots of 4 litres containing a mix of sterilized unfertilized peat (40% Baltic blond peat 0-10 mm, 60% Brown peat 0-10 mm, 75 g oligo elements, pH CaCl_2_ 5.3 – 5.8, Stender, France) and dried clay beads (reference: 17831050 BILLE ARGILE LIA 8/16, Soufflet, France) 1:1/v:v. For experimental design and the different steps of growing and harvesting of the experiment in 2020 and 2021 see Supplementary Figure 1 and 2 respectively. For meteorological conditions within the greenhouse during 2020 see Supplementary Figure 3. For each condition, 9-10 plants were used. Three inoculations with spores of *R. irregularis* were performed as follow: The first inoculation with 500 spores/plant at sowing, second inoculation 1 week after sowing with 200 spores/plant and a third inoculation 2 weeks after sowing with 200 spores/plant. Control plants were treated with distilled water. Plants were grown under three different nitrogen regimes: low N (LN, 1mM NO_3_^−^), medium N (MN, 5 mM NO_3_^−^) and high N (HN, 10 mM NO_3_^−^). The specific composition of these regimes is provided in Supplementary Table 2. For each N regime, plants were irrigated for the first three weeks with the solution corresponding to the chosen N regime with low P (named P-) containing 10 µM phosphate in order to trigger AMF colonization. Then, at dates indicated in Supplementary Figure 1 and 2, irrigation was changed to the same N level but 300 µM P (named P+, Supplementary Table 2). Increasing P level in the irrigation solution was necessary to allow plants reach the maturity stage. To determine ^15^N uptake, ^15^N labelling was performed at indicated dates (Supplementary Figure 1). 100 mL of 45 mM KNO_3_ solution enriched at 2% ^15^N was applied to each plant in order to provide 1,25mg of 15_N._

### Harvesting procedures

Plants grown in 2020 were used for the phenotyping, metabolomics, RNA-seq and RT-qPCR experiments. Plants grown in 2021 were used to monitor fungal colonization and RT-qPCR experiments. Depending on the experiment and the biological question asked, one, two or three developmental stages were chosen for tissue harvesting. The first one is the vegetative stage (VS) corresponding to the 7– to 8-leaf stage ^36^ is an indicator of a highly metabolically and photosynthetically active leaf tissue. The second stage corresponds to 15 days after the tasselling (VT+15) and plant is in the early phase of silking (data not shown). This stage was chosen because it allows root harvesting at a reproductive stage without compromising kernel production. The third stage is the maturity stage (MS) and corresponds to plants displaying mature ears. Dates of harvesting for VS, VT+15 stage and MS differed depending on the condition and the year of experiment (Supplementary Figure 1 and 2). For leaf tissues, harvesting consisted in cutting a piece of 1 cm x 10 cm of leaf mesophyll along the midrib from the 6^th^ fully emerged leaf for the VS. For VT+15 stage and MS, a piece of 1 cm x 10 cm leaf surface from the leaf below the ear was harvested. Roots were harvested at VS, VT+15 stage or MS depending on the type of experiment. For each treatment/condition, roots were collected from 5 plants. Thin secondary roots were harvested from the surrounding root system, close to the pot borders. They were washed with distilled water, then frozen in liquid nitrogen or stored in distilled water. Following root harvesting, the plants grown in 2020 grew until maturity.

### Quantification of yield, biomass, chlorophyll, C, N, ^15^N and P

Plant organs were harvested then oven dried at 80°C for 3 to 5 days then ground to fine powder. Grain, biomass of the aerial parts including stalks, leaves, and cobs were weighed after drying. The upper regions of the root system was harvested. The percentage of C and N were quantified using the FLASH 2000 Organic Elemental Analyzer (Thermo Fisher Scientific Villebon, France) and the ^15^N enrichment using the Delta V Advantage isotope ratio mass spectrometer (Thermo Fisher Scientific, France) as described in ^37^. Supplementary Table 3 indicates the calculations of different traits with C and N content. A% (Atom percent) represents the ^15^N enrichment in the sample [^15^N/(total N)]. Since the natural ^15^N abundance in N unlabelled samples was 0.3663 (A% control), specific enrichments due to the ^15^N uptake were calculated as E% = (A% – 0.3363).The P concentration was measured using the molybdate blue method on a Shimadzu UV-160 spectrophotometer (Shimadzu Biotech) after acid digestion ^38^.

### Quantification of root colonization level by AMF structures

Thin secondary roots were harvested as indicated above and carefully washed in distilled water. They were incubated in 10% KOH overnight at room temperature. They were then rinsed in distilled water and incubated in 5% black ink (Sheaffer Skrip Bottled Ink, France) solution for 10 min at 70°C. Washing was performed by incubating roots in 8% acetic acid for 25 min. For the quantification of % of AM colonization (total, arbuscules, vesicles) 50 pieces of 1 cm length each were mounted on a slide for observation of AM structures under light microscope using the slide method ^39^. Quantifications were performed on roots collected from 5 plants for each condition.

### RNA extraction for RT qPCR and RNAseq

RNA extraction was performed with TRIzol (Thermo Fisher Scientific, USA) protocol. Fresh tissue of leaves or roots (100 mg) were frozen in liquid nitrogen then ground to fine powder that was homogenized in 1 mL of TRIzol. RNA-extraction was performed as described in ^34^ and was used for RT-qPCR and RNA-seq.

### cDNA synthesis and RT-PCR

It was performed as indicated in Decouard et al. ^37^. Primer sequences are described in Supplementary table 4.

### RNA-seq and Bioinformatic Analysis

RNA-seq libraries were constructed by the POPS platform (IPS2) using the Illumina Stranded mRNA Prep kit (Illumina®, California, U.S.A.) according to the supplier’s instructions. Libraries were sequenced by the Genoscope (Évry-Courcouronnes, France) on a Novaseq 6000 instrument (Illumina) with 150 base paired-end sequences. RNA-seq raw data cleaning was made as described in ^34^. Statistical RNA-seq analyses are described in Supplementary File 1. Filtered reads were then mapped and counted using STAR (v2.7.3a, ^40^) with the following parameters –-alignIntronMin 5 –-alignIntronMax 60000 –-outSAMprimaryFlag AllBestScore –-outFilterMultimapScoreRange 0 –-outFilterMultimapNmax 20 on a combined reference genome made by concatenating the files of the *Zea mays* B73 (V4.49) and *Rhizophagus irregularis* (V2) reference genomes and the associated GTF annotation files (also combined). Between 75.3 and 96.1 % of the reads were uniquely mapped (median = 91.9%). Between 68 and 88 % of the reads were associated to annotated genes (median = 85.3%).

### Metabolomic analyses

All extraction steps were performed in 2 mL Safelock Eppendorf tubes. The ground dried samples (5 mg DW) were resuspended in 1 mL of frozen (–20°C) Water:Acetonitrile:Isopropanol (2:3:3) containing Ribitol at 4 mg/L and extracted for 10 min at 4°C with shaking at 1500 rpm in an Eppendorf Thermomixer. Insoluble material was removed by centrifugation at 13500 rpm for 10 min. 700 µL were collected and 70 μL of myristic acid d27 at 0.3 g/L was added as an internal standard for retention time locking. After centrifugation, aliquots of each extract (100 µL) were dried for 4 h at 35 °C in a Speed-Vac and stored at –80°C. Two Blank tubes underwent the same steps as the samples.

Sample derivatization and analysis: All steps of GC-MS analyses were done as described in Fiehn et al.^41,42^. Agilent literature PUB: 5989-8310EN (Agilent G1676AA Agilent Fiehn GC/MS Metabolomics RTL Library User Guide. June 2008. Agilent P/N: G1676-90000, Agilent Fiehn GC/MS Metabolomics RTL Library Product Note. May 2008). Samples were taken out of –80°C, warmed 15 min before opening and speed-vac dried again for 1 hour at 35 °C before adding 10 µl of 20 mg/mL methoxyamine in pyridine to the samples and the reaction was performed for 90 min at 30°C under continuous shaking in an Eppendorf thermomixer. 90 µL of N-methyl-N-trimethylsilyl-trifluoroacetamide (MSTFA) (REGIS TECHNOLOGIES 1-270590-200 –10×1g) were then added and the reaction continued for 30 min at 37°C. After cooling, 90 µL were transferred to an Agilent vial for injection.

4 hours after derivatization 1 µL of sample was injected in splitless mode on an Agilent 7890B gas chromatograph coupled to an Agilent 5977A mass spectrometer. The column was an ZORBAX DB5-MS from Agilent (30 m with 10 m integraguard column – ref 122-5532G). An injection in split mode with a ratio of 1:30 was systematically performed for saturated quantification compounds. Oven temperature ramp was 60 °C for 1 min then 10 °C/min to 325 °C for 10 min. Helium constant flow was 1.1 mL/min. Temperatures were as follows: injector: 250°C, transfer line: 290°C, source: 230 °C and quadripole 150 °C. The quadrupole mass spectrometer was switched on after a 5.90 min solvent delay time, scanning from 50-600 µ. Absolute retention times were locked to the internal standard d27-myristic acid using the RTL system provided in Agilent’s Masshunter software. Retention time locking reduces run-to-run retention time variation. Samples were randomized. A fatty acid methyl esters mix (C8, C9, C10, C12, C14, C16, C18, C20, C22, C24, C26, C28, C30) was injected in the middle of the queue for external RI calibration.

Data processing and quantification: Raw Agilent datafiles were analyzed with AMDIS http://chemdata.nist.gov/mass-spc/amdis/. The Agilent Fiehn GC/MS Metabolomics RTL Library (version June 2008) was employed for metabolite identifications. This library is the most comprehensive library of metabolite GC/MS spectra that is commercially available. It contains searchable GC/MS EI spectra and retention time indexes from approximately 700 common metabolites. Peak areas determined with the Masshunter Quantitative Analysis (Agilent) in splitless and split 30 modes. Because automated peak integration was occasionally erroneous, integration was verified manually for each compound in all analyses. Resulting areas were compiled in one single Excel file for comparison. Peak areas were normalized to Ribitol and Dry Weight. Metabolite contents are expressed in arbitrary units (semi-quantitative determination).

### Amino acid quantification by OPA-HPLC

Extraction procedure was identical that one described for metabolomics. Here 200µL of the extraction solution was taken and dried in a Speed-Vac evaporator for 2 h at 30°C before adding 1,3 mL of H_2_0 milliQ. The o-phtaldialdéhyde (OPA) reagent was made 48 h before first use by dissolving OPA at 12mg/mL in 200 µL of methanol and adding 1.8 mL of sodium borate 0.5M (pH 9.5) and 40 µL of 2-mercaptoethanol. The reagent was filtered into an autosampler vial and used for up to 3 days. Precolumn derivatization was performed in the injection loop by automated mixing of 10 µL sample and 10 µL OPA reagent, followed by a delay of 2 min prior to injection. The HPLC (Alliance Waters 2695, Waters Corporation, Massachusetts, USA) fitted with a fluo detector (Multi λ Fluorescence Detector 2475) was used. Compounds were measured at λ excitation of 340 nm and λ emission of 455 nm. The chromatographic separation was performed with Symmetry C18 column (3.5µm, 4.6*150 mm) by gradient elution at 40 °C using buffer A (20% methanol, 80% sodium acetate, 1% tetrahydrofuran, pH 5.9) and buffer B (80% methanol, 20% sodium acetate, pH 5.9). Buffer flow rate was 0.8 mL/min throughout and total run time per injection was 42 min. The chromatography data were analysed by Empower software. Peak identity was confirmed by co-elution with authentic standards. Concentration was calculated with calibration curve using peak area of compound of interest.

### Nucleotide quantification LC–TOF

Nucleotide quantification as performed on frozen leaves of the same maize plants used for metabolome and RNAseq. Extraction and quantification were performed as described in ^43^.

### Statistical analyses

All statistical analyses were performed with RStudio 2023.06.0+421 “Mountain Hydrangea” under the free software environment R version 4.3.1 (Beagle Scouts). All Anova were implemented with the “aov” R package. All heatmaps were generated with the estimated marginal means that were calculated using the R package “emmeans”. Heatmap for physio-agronomical traits was generated with ‘pheatmap’ 1.0.12 package (Date 2019-01-04. Author Raivo Kolde) by implementing the ward.D2 hierarchical clustering method ^44^. Heatmap for physio-agronomical traits was generated with the ‘pheatmap’ R package 1.0.12 Date 2019-01-04 (Author Raivo Kolde). Heatmaps for metabolomics, including the interactive ones, were generated with “heatmaply” R package ^45^.

Metabolomics data were analyzed with the R package MultiVarSel ^46^ which is the implementation of the variable selection approach in the multivariate linear model proposed in ^47^ theoretically analysed in ^46^ and applied in several contexts of omics data analyses such as ^48,49^. The list of R packages used in this study *sbm: Stochastic Blockmodels*. R package version 0.4.5-9000, <https://grosssbm.github.io/sbm/>. Kolde R (2019). _pheatmap: Pretty Heatmaps_. R package version 1.0.12, <https://CRAN.R-project.org/package=pheatmap>. Lenth R (2023). _emmeans: Estimated Marginal Means, aka Least-Squares Means_. R package version 1.8.6, <https://CRAN.R-project.org/package=emmeans>. Perrot-Dockès M, Lévy-Leduc C, Chiquet J. R package MultiVarSel available from the Comprehensive R Archive Network, 2017. <https://CRAN.R-project.org/package=MultiVarSel>.

### Genome-Scale Metabolic Modelling (GSM) and Transcriptomics Data Integration

To investigate the metabolism of maize from systems level, we used *i*ZMA6517 as the template model ^34^. The model is specific to B73 genotype and consists of organ-specific GSMs of root, stalk, leaf, and kernel. All these individual organ GSMs are connected through a common phloem pool. The model has been described in our earlier work ^34^. The model was contextualized with HN Control, HN Rhizo, LN Control, and LN Rhizo using EXTREAM algorithm ^34^. The upper and lower limits of the reaction fluxes for the mentioned conditions are presented in the Supplementary Table 5. The General Algebraic Modeling System (GAMS) version 24.7.4 with IBM CPLEX solver was used to run FBA and FVA, E-Flux, and the FSA algorithm on the model. Each of the algorithms was scripted in GAMS and then run on a Linux-based high-performance cluster computing system at the University of Nebraska-Lincoln. Instructions for running files in GAMS through NEOS server have been published ^50^. All the necessary files for COBRApy users can also be accessed in the GitHub depository (https://github.com/ssbio/iZMA6517).

### Metabolic Bottleneck Analysis (MBA)

To determine the metabolic bottlenecks in each of HN Control, HN Rhizo, LN Control, and LN Rhizo conditions, we used Metabolic Bottleneck Analysis (MBA) ^34^.. MBA expands the flux space of each reaction separately and assesses its impact on the whole plant biomass growth rate. Further details on MBA can be accessed in the original publications. MBA pinpoints plant-wide metabolic bottlenecks, which were further analyzed for biological insights.

## Results

### The impact of *Rhizophagus irregularis* on maize growth and yield is strongly dependent on the level of N nutrition

To investigate the diversity of the molecular and physiological mechanisms involved in maize interaction with AMF in terms of N nutrition we used two genetically distant AMF-responsive maize lines, B73 and Lo3. B73 is known be responsive to AMF, resulting in increased biomass ^25,51^. Lo3 genotype was selected due to its enhanced biomass response following inoculation with *R. irregularis* in two rounds of selection (data not shown). In order to find at which level of N fertilization the two lines benefit most from the fungal inoculation, these were grown in a greenhouse under low N (LN), medium N (MN) or high N (HN) corresponding to 1 mM NO_3_^−^, 5 mM NO_3_ ^−^ and 10 mM NO_3_ ^−^ in the nutrient solution respectively. The impact of *R. irregularis* inoculation on a panel of maize physio-agronomic traits was monitored (Supplementary table 3). *R. irregularis* inoculation resulted in an increase in the shoot biomass production (Aerial Parts, AP) and grain yield under LN in both lines (Figure 1A, 1B), and under MN for B73. A negative impact of *R. irregularis* inoculation was observed on B73 grain yield when plants were grown under HN (Figure 1B). Interestingly, yield of mycorrhizal plants grown under LN was similar to yield of non-inoculated plants fed with 5 times more NO_3_^−^ (MN). NUE increased in the AP only in LN conditions for both inoculated lines compared to controls (Figure 1C). An increase in NUE for grain was observed under LN and MN fertilization conditions only in B73 (Figure 1D). Fungal inoculation resulted in a significant increase in the N content of aerial parts under MN for B73 and under HN for Lo3 indicating that these lines are able to store higher levels of N in the vegetative parts upon association with the AMF even under ample N (Supplementary Figure 4A, B, C). To determine whether N uptake is modified during the symbiotic interaction, plants were labelled with a pulse of ^15^N provided at the vegetative stage. The total quantity of ^15^N in aerial parts and grains measured at maturity remained unchanged whatever the treatment (Supplementary Figure 5D and E), except for the AP of Lo3 under HN. In inoculated plants, an increase in the total phosphorus content was observed mainly under HN conditions (Supplementary Figure 4E, F, G). Hierarchical clustering analysis (HCA) was performed using the physio-agronomic traits listed in Supplementary Table 3 (Figure 1 E). The HCA shows that the N supply is the major driver of all traits. All the LN plants were grouped in a cluster where values for the traits related to AP and roots were higher. On the contrary, in cluster B, traits related mainly to grains were higher. In cluster B, genotypes were distributed into two subgroups B1 and B2. Interestingly, in cluster A, traits related to grains exhibited higher values in the inoculated plants compared to the control indicating that *R. irregularis* allowed plants to behave more similarly to group B plants, albeit still under N stress deficiency. These data show that *R. irregularis* inoculation can mitigate the impact of N limitation on physio-agronomic traits.

**Figure 1:**
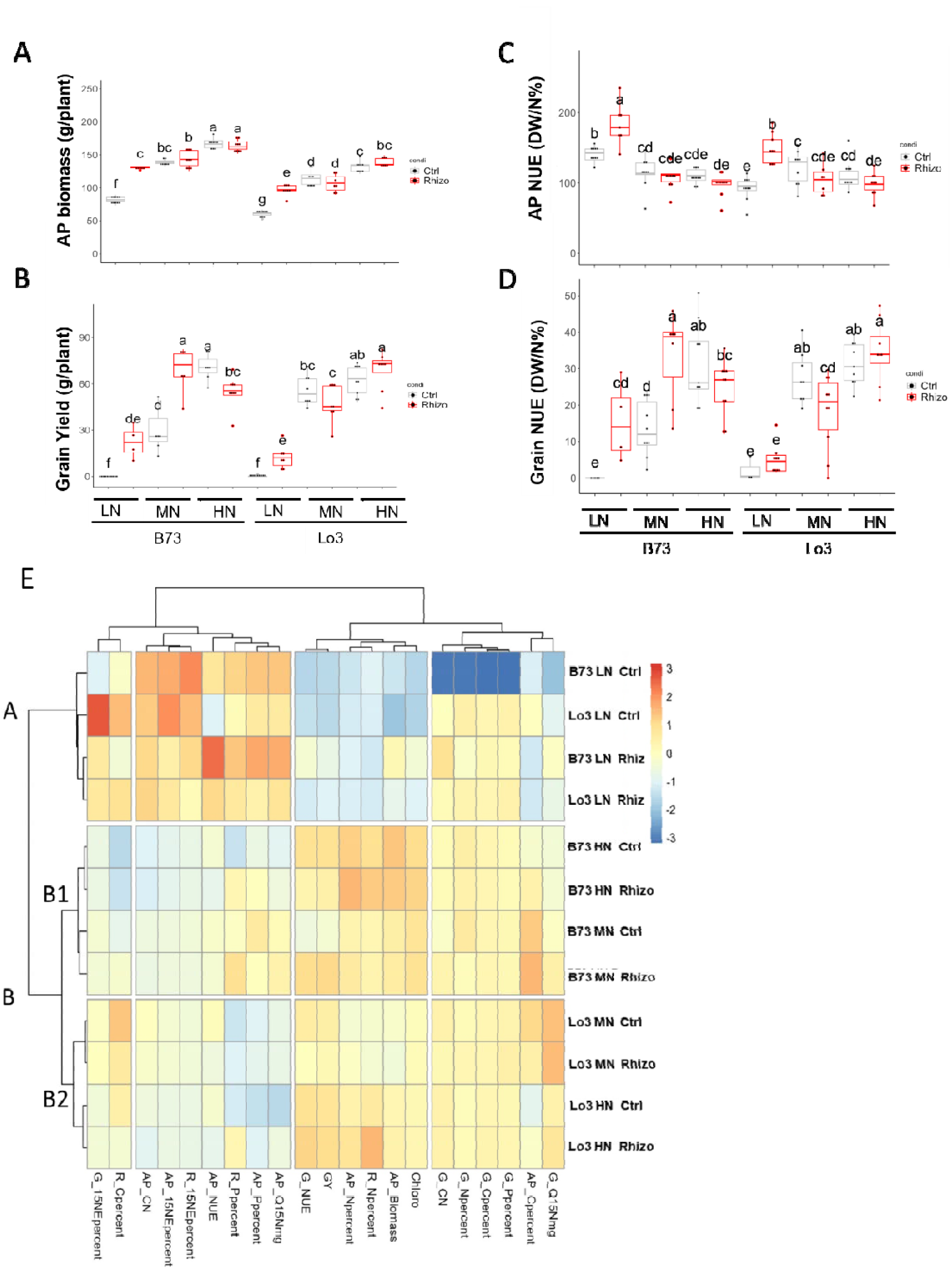
Impact of N supply on maize responses ta *R*. irregularis for biomass and N content shows more pronounced beneflclal AMF effects undar LN. **A and, B**: grain yield and aerial part (AP) biomass. C and D: Nitrogen Use Efficiency (NUE) in grains and in NP. Gtrl: Mock non inoculated plants, Rhizo: inoculatec with *R*. irregularis. Boxplots depict the interquartile range, with the median demarcated by a horizontal line in the micdle, upper, and lower part of the box, representing the first and third quartiles and the range of data distribution. Dots overlaying the boxplots n1present repentant plants, each corresponding to a single observation In the dadaset (N=7 to 10 plants). Different letters indicate significant differences betweetn groups identified through one-way ANOVA testing followed by Tukey’s Honestly Significant Difference (HSD) post hoc analysis (p < 0.05). An asterisk indicates a significant difference between control and inoculated plants, determined by a t-test (p < 0.05). **E**: Hlerarchical Clustering Analysis was performed folowing Ward method of clustering. Indicated traits are measured in Aerial parts (AP). roots (R) or grains (G). Cpercen;: Percentage of C. Npercent: percentage of N. C/N: ratio. 15NEpercent: percentage of ^15^N. Ppercent: percentage of P. Q15Nmg: quantity of 15N in organ per plant. NUE: Nitrogen Use Efficiency, GY: Grain Yield our plant. The color scale Incloates If the tralt is higher (oranga) or lower (blue) In the Indicated condition, with respect to the others. Details for the formulas used to calculate these traits are indicated in Supplementary Table 3.

### Marker genes for N uptake and assimilation are differentially regulated in response to AMF inoculation and N supply

The finding that several NUE traits, notably those related to NO_3_ ^−^ uptake were modified by AMF treatment prompted us to investigate further which molecular components of symbiosis and N transport were involved at the root level. Roots were harvested at the VT+15 stage, corresponding to 15 days after tasselling (see materials and methods) that was identified as the most relevant stage to investigate maize-AMF interaction (Supplementary Figure 6). By monitoring the level of *R. irregularis* constitutive transcript *RiLSU,* a constitutive *R. irregularis* transcript, we found that high N input led to increased fungal biomass in roots (Figure 2A). Then, the level of transcripts for the plant genes *ZmAMT3.1*, *ZmPHT6.1; ZmHA1* and *ZmNPF4.5* markers of the symbiotic association and representative of NH_4_^+^, phosphorus and NO_3_^−^ transport respectively was quantified by qRT-PCR ^13,27,52,53^.

**Figure 2:**
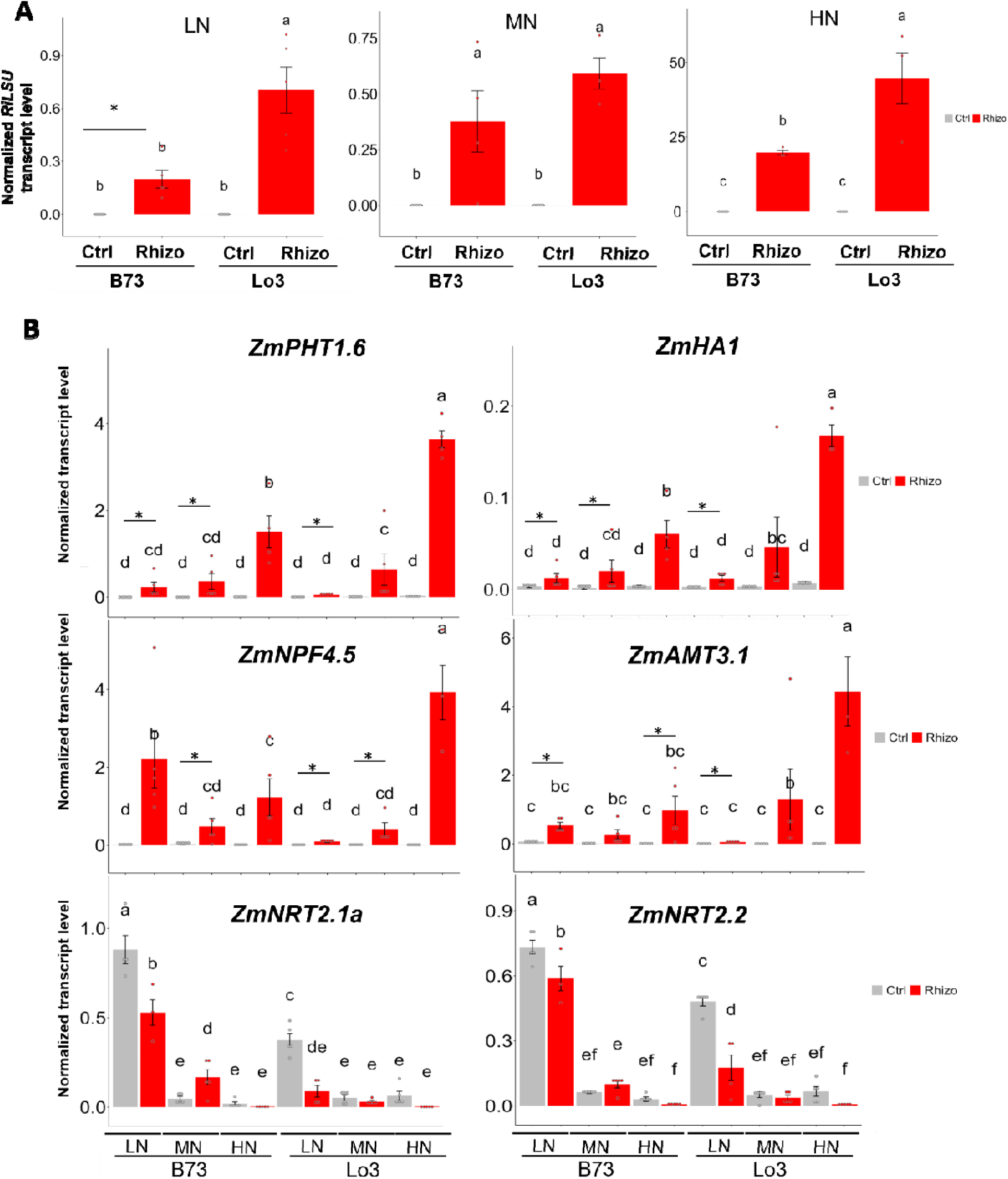
Impact of N supply on expression of genes Involved In N end P transport In roots suggests that AM symbiosis alleviates N deficiency stress. Plente were grown end harvested In 2020 as Indicated In materiale and methods under law nitrate (LN, 1mM NO_3_-), moderate nitrate (MN, SmM NO_3_-) or high nitrate (HN, 10mV NQg). Rnate were harvested at VT+15 stage. A: Quantification of *R. Imgularia* In mate by qRT-PCR amplificatIan of the *RILSU* constitutive fungal transcript normalized with maize actin transcripta. B: Normalized transcript levels with actin as a reference for Indicated genes monitored by qRT-PCR. Ctrl: Mock non Inoculated plente, Rhlzo: Inoculated with *R. imgulariB* Bar plots represent the mesne, accompanied by dots for Individual observations overlaid on the bars, with each dot corresponding to a single observation In the dataset (N=5). Different letters Indicate significant differences In gene expression levels Identified through one-way ANOVA analysis followed by Tukey’s honestly significant difference (HSD) test. The black bars represent standard errors. An asterisk Indicates a statistically significant difference between mock and *R.* irraguferfs-lnoculoted plants, determined by a bra-somple t-fest (p < 0.05).

The expression level of the four well-known symbiotic markers mentioned previously, *ZmPHT1.6*, *ZmNPF4.5*, *ZmHA1*, a proton pump, and *ZmAMT3.1* is up-regulated in response to the fungus in both genotypes and under all N supply conditions (Figure 2B). It is interesting to notice that the higher the N nutrition level, the higher the expression of these markers (Figure 2B). To know more about direct N transport genes in those conditions, we investigated expression levels of two maker genes of N starvation stress *(ZmNRT2.1a, and ZmNRT2.2)* encoding high affinity transporters, known to be upregulated under N deficiency ^2,54–56^ (Figure 2B). As expected, the level of N supply had a drastic effect in both maize genotypes on transcripts levels for the nitrate transporters *ZmNRT2.1a* and *ZmNRT2.2* which were more highly expressed in LN plants compared to MN and HN plants. Interestingly, the inoculation with *R. irregularis* on LN plants resulted in decreased levels of the *ZmNRT2.1a* and *ZmNRT2.2* transcripts. These data indicate that the inoculation of N starved plants with the fungus mimics an additional N supply.

### Leaf and root transcriptome reprogramming in response to *R. irregularis* colonization integrates a genotype and N-availability effects

To further characterize the biological processes involved in maize response to *R. irregularis* colonization as a function of N fertilization, a whole genome transcriptomics approach was conducted in leaves and a dual RNA-seq in roots of plants grown under HN and LN conditions. The MN condition was considered as equivalent to HN and was not further investigated. The leaf below the ear was chosen for providing a good proxy of the sink-to-source transition during grain filling ^57^. First, we defined the differentially expressed genes (DEGs) as those exhibiting a fold change of the transcript levels in inoculated plants compared to non-inoculated plants that was higher than 1.5 or lower than –1.5 (*p*<0.05). Details for the numbers of DEGs depending on organ, N nutrition level and genotype are shown in Figure 3. GO enrichment analysis revealed that for those common up-regulated genes in roots, the biological processes overrepresented were those related to proteolysis, lipid metabolism, ethylene response and membrane transport (Supplementary Figure 7). The common down regulated genes were represented by five main functional categories including cell wall biogenesis, metal homeostasis, phosphate transport, stress responses and lipid metabolism (Supplementary Figure 7). Specific features of B73 root transcriptome consist mainly on up-regulation of genes related to immunity (both under LN and HN). Specific features of the root Lo3 transcriptome resides mainly on up-regulation of autophagy related genes (under LN) and terpenoid metabolism genes, and down regulation of genes related to DNA metabolism and replication (both under LN and HN) (Supplementary Figure 7). We performed the same data analysis in leaves (Figure 3, Supplementary Figure 7). Specific features of B73 leaf transcriptome consist mainly on translation and auxin signalling genes (under LN), and redox reactions, carbohydrate transport and defence related genes (under HN). Specific features of the leaf Lo3 transcriptome reside in photosynthesis, cell wall remodelling, cytoskeleton remodelling, stress response, signalling, while in the down-regulated genes are related to gamma-amino butyric acid (GABA) metabolism and pyrimidine biosynthesis (under both LN and HN). Taken together, these data show that leaf and root transcriptomic responses to the AMF differ depending on maize genotype and N supply.

**Figure 3:**
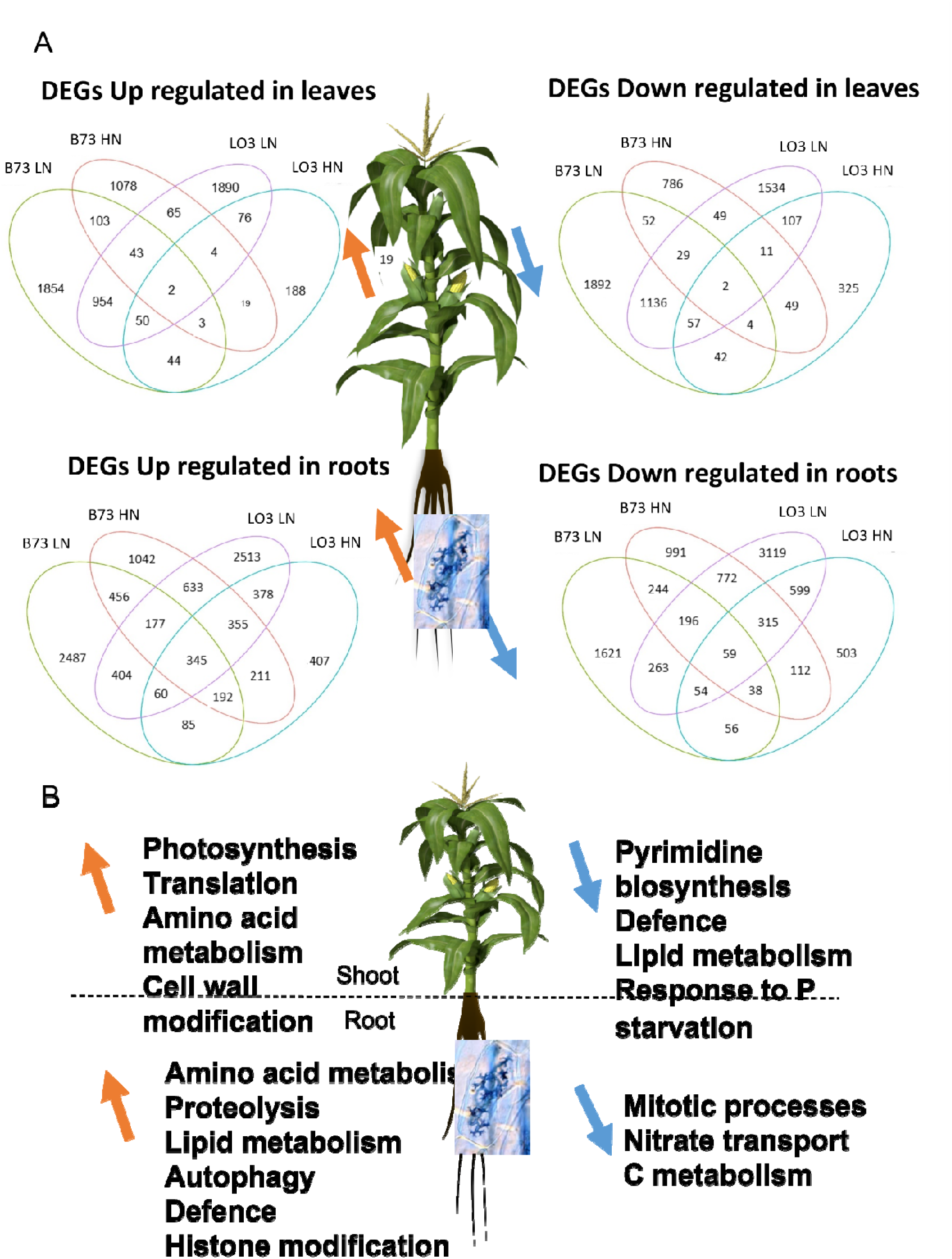
RNAssq data showing the overlap between main DEGs and functional GO enrichment categoriae. A: Ganes were selected so that their fold change (inoculated varsus control) was > to 1.5 and statistically significent in the 3 teplicatae. В73 LN end B73 HN represent DEG genae up or down regulated In Inoculated plente compand to control plants In LN or HN respectively; LO3 LN and LO3 HN rapraaant DEG genes up or down regulated n Inoculated plants compered to control plente In LN or HN reeoectvely. B: mein GO functional categories in DEGs down h Inoculated venue control (on the right with blue arrows] and DEG down (on the left with orange arrows) hi Inoculated versus control.

### Leaf and root metabolomics profiles in response to *R. irregularis* are shaped by N nutrition

To further characterize the metabolic responses of the two maize lines to *R. irregularis* inoculation, a primary metabolome profiling was performed on the same leaf and root samples as those studied in the previous section. Again, plant physio-agronomical traits and metabolomics are paired data. Metabolic profiles showed that plant organ and N supply had the strongest impact on leaf and root metabolite composition (Supplementary Figures 8A and B). AMF inoculation and maize genotype had globally weaker impacts on metabolite profiles (Figures 4 A and B and supplementary Figure 8C and D). In leaf samples, 132 water soluble metabolites were detected of which 103 were significantly different in at least on condition after ANOVA statistical analysis (*P* < 0.05) followed by a Bonferroni post-hoc test and correction. In roots, there were 120 water-soluble metabolites of which 117 were found to be significantly different in at least one condition. The identity of metabolites can be found on the interactive heatmaps (Supplementary interactive Files 1 and 2).

**Figure 4:**
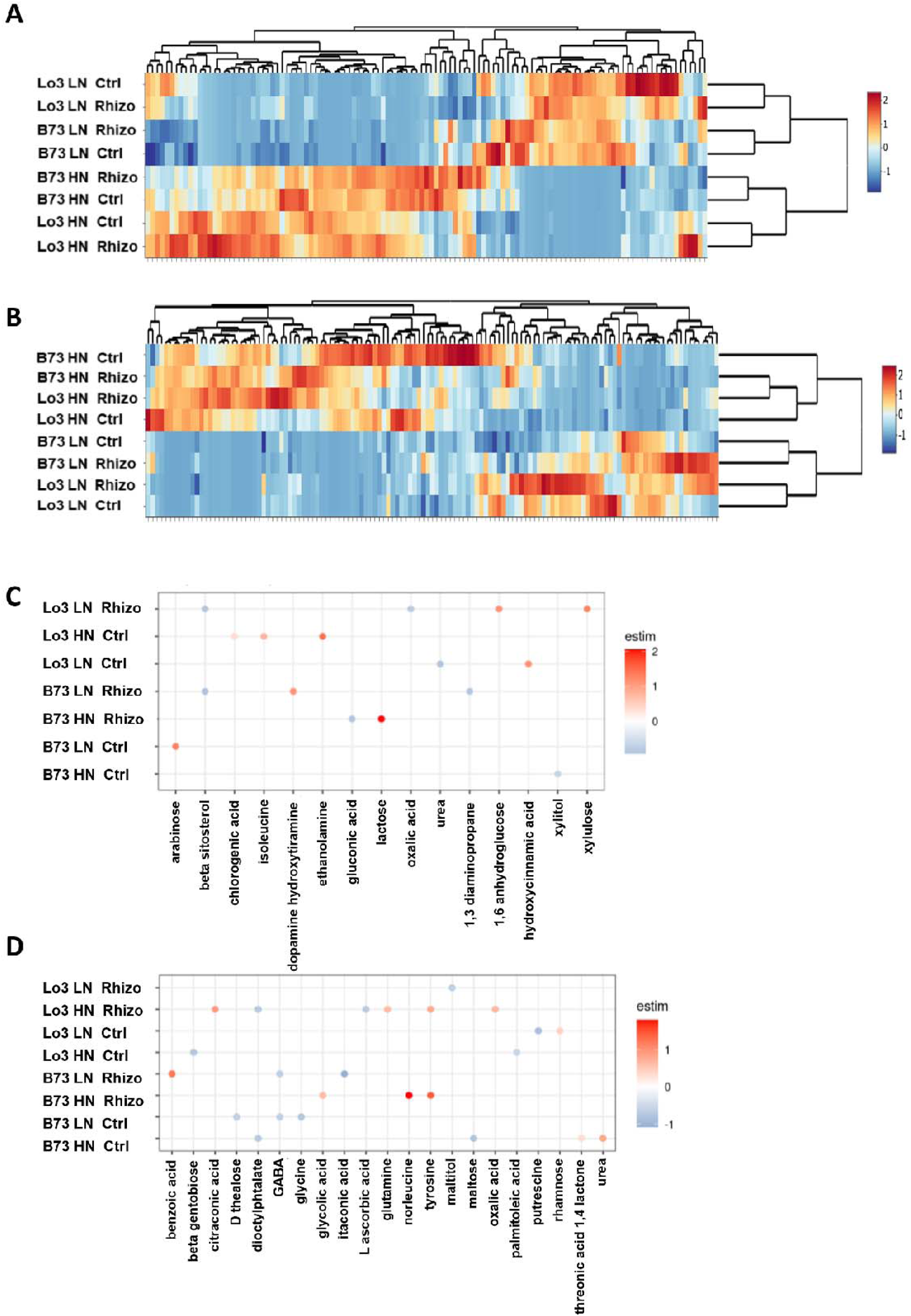
M**e**tabolomic **analyses of leaves and roots reveals key metabolites specific to symbiosis under LN or HN nutrition.** Plants were grown and harvested in 2020 as indicated in materials and methods under low nitrate (LN, 1mM NO_3_-) or high nitrate (HN, 10mM NO_3_-). Leaves and roots were harvested at VT+15 stage. Ctrl: Mock non inoculated control, Rhizo: inoculated with *R. irregularis,* Metabolites were monitored by GS/MS. **A** and **B:** HCA heatmap showing metabolomic profiles of leaves and roots of inoculated or control plants under LN or HN in leaves and roots respectively. **C** and **D**: Analyses of metabolomics data conducted with the R package MultiVarSel with a threshold of 0.884 showing the metabolites on which the genotype, the N nutrition of the *R. irregularis* inoculation have an impact. Leaves, **C;** roots, **D.** The color scale indicates if the metabolites are accumulated (red) or down accumulated (blue) in the indicated conditions with respect to the others.

In roots, fungal inoculation triggered accumulation of lipids such as trans-13-octadecenoic acid and palmitoleic in both lines Supplementary Figure 9 A. The accumulation of these two lipids was much higher under HN than under LN. We observed that in inoculated roots, there was a global decrease in sucrose, glucose and fructose levels Supplementary Figure 9 B. Several metabolites linked to N metabolism were differentially accumulated in inoculated roots Supplementary Figure 9 D. For example, Glu, Arg, citrulline, Asn. Some metabolites related to immunity were accumulated differentially in response to the fungus. For instance, shikimic acid was accumulated in LN roots of inoculated B73 plants compared to controls Supplementary Figure 9 C. In leaves, sucrose showed a significant decrease in HN in both maize lines (Supplementary Figure 10). Gln and Asn accumulated in Lo3 leaves under HN in response to *R. irregularis*.

To identify metabolites representative of the plant response to *R. irregularis* under each growth condition, metabolites for which abundance is significantly correlated with specific treatment conditions were identified by using a variable selection approach in a multivariate linear model conducted with the MultiVarSel approach (see materials and methods, Figure 4 C and D). For example, we found that under LN, xylulose, was correlated with leaves of Lo3 inoculated plants (Figure 4C). In roots, benzoic acid was accumulated in inoculated B73 roots of plants grown under LN (Figure 4D). These data indicate that specific metabolites accumulation is correlated with particular conditions depending on the plant genotype and the N supply in response to *R. irregularis*.

### Multi-organ metabolic modelling reveals key metabolic processes triggered by *R. irregularis* allowing maize to mitigate N limitation stress

To identify key metabolic reactions involved in yield and biomass reshaping in response to *R. irregularis* symbiosis, we integrated the transcriptomic data in a recently reconstructed multi-organ genome scale modelling (GSM) iZMA6517 using EXTREAM ^34^. This was followed by a metabolic bottleneck analysis (MBA) ^34^ to pinpoint metabolic bottlenecks for biomass growth rate. First, the B73 leaf and root transcriptomic data were used as input to EXTREAM to contextualize iZMA6517. EXTREAM equally distributes transcript levels for each gene based on the number of cognate biochemical reactions and contextualize the GSM. The in silico analysis predicted that growth rates were increased under LN in response to *R. irregularis* inoculation compared to non-inoculated plants, which is expected to lead to grain yield and biomass increase. While under HN, a negative impact on maize growth rate was predicted (Figure 5A). In particular, the model predicted absence of grain production in control of LN B73 plants. Strikingly, these model predictions agree with the observed yield and biomass experimental data in Figure 1, confirming the predictive accuracy of the iZMA6517. When applied to the Lo3 genotype leaf and root transcriptomic data, iZMA6517 predicted a slight increase in grain yield and leaf biomass under LN in Lo3 inoculated plants. The model was better fitting the experimental B73 phenotype in terms of biomass compared to the Lo3 experimental data.

**Figure 5.**
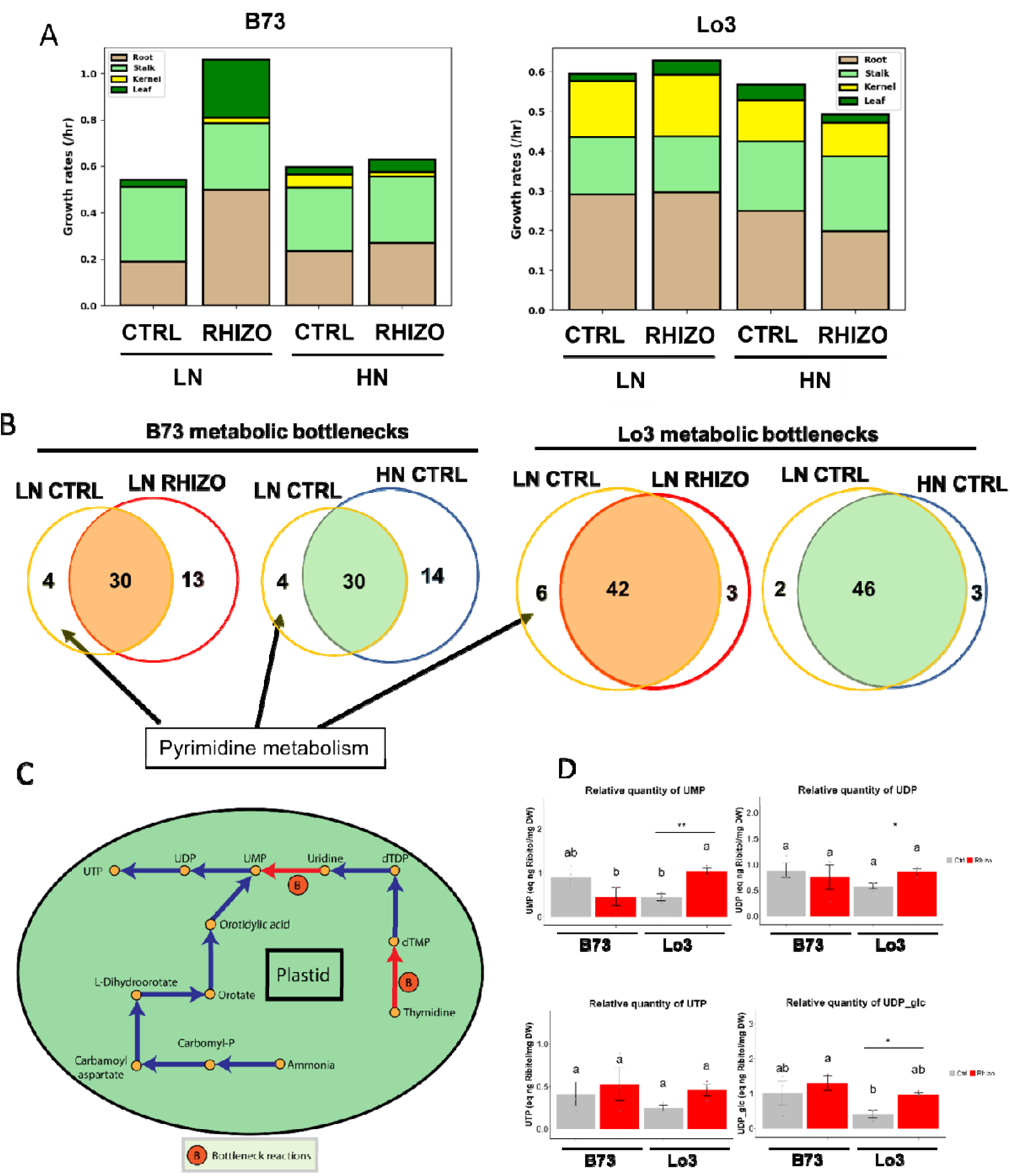
: Genoma baaed metabolic modeling reveals essential impact of ДМҒ on pyrimidine matsboliam. A: RNAaeq data were used to predict different organ growth rates In the B73 and Lo3 maize lines. EXTREAM algorithm predicted higher yield under LM In В73 and Lo3. B: Venn diagrams showing the metabolic bottleneck» under Indicated conditions. The debottlenecked reactions from the LN CTRL condition ere those on the left yellow circle outside the intersect on. LN CTRL (yellow circle). HN CTRL (blue circle). LN RHIZO (red circle). The list of the metabolic bottlenecks Is on Tables 1 and 2. C; Metabolic reactions related to the pyrimidine pathway that ere predicted to be dcbettloncekcd by *R. Irregularia* Inoculation and by HN tmetmarrt shown In B. All the reactions shown hare are In the pbntld comoartmanl. D: Quantification of pyrimidines In leaves of the same plants from the 2020 experiment, used tor RNAaeq and metabolomlcs. The nucleotides were extracted then quert tied by LC-TOF according to materials and methods. N”3 pools of leaves harvested from 2 to 3 plants. CTRL Mock non Inoculated, RHIZO; Inoculated with *R. Irregularia.* Different letters Indicate significant differences In nucleotide levels Identified through one-way ANOVA analysis followed by Tukey*e honestly significant difference (HSD) test. The black bars represent standard errors. Ona or two asterisks Indicates a statistically significent difference between mock and R. *Irragularia-Inoculated* plants, determined by a two-sample t-test p < 0.05 or p<0.01 respectively

This result prompted us to determine the metabolic processes that could explain the improved yield under LN B73 plants and LN Lo3 in response to *R. irregularis* inoculation by identifying the potential metabolic bottlenecks relieved by the AMF under limiting N using MBA (see materials and methods). MBA expands the flux space of each reaction individually and assesses the impact of the expanded flux space on the biomass growth rate. For this purpose, we compared the bottlenecks identified in LN control plants to the bottlenecks identified in inoculated LN plants (Figure 5B; Tables 1 and 2). Four metabolic bottlenecks relieved by *R. irregularis* inoculation were identified (Table 1). Interestingly 13 new bottlenecks were emerged following *R. irregularis* inoculation. On the other hand, we found that four metabolic bottlenecks that were present in LN control plants were relieved by N supply in the HN plants. There were three metabolic bottlenecks relieved by both HN supply conditions and *R. irregularis* inoculated condition (photosynthesis, purine and pyrimidine metabolism, aromatic acid metabolism). The same analysis was conducted in Lo3 maize genotype. There were 6 metabolic bottlenecks relieved by *R. irregularis* inoculation under LN conditions (Table 2). These bottlenecks are involved in Pyrimidine metabolism, Purine metabolism, and Calvin-Benson-Bassham cycle. These data suggest that in both maize lines, *R. irregularis* inoculation triggers modifications in pyrimidine metabolism. More specifically, the model predicts that the limiting reactions that are relieved by symbiosis should result in an accumulation of UMP, UDP and UTP in leaves under LN. Thus, the levels of UMP, UDP, UDP-glc and UTP were monitored in leaves under LN and HN (Figure 5D and Supplementary Figure 12). Remarkably, we found an increase in UMP, UDP and UDP-glc levels in leaves of plants inoculated with the AMF under LN only in the Lo3 genotype, thus confirming MBA predictions. Together these data indicate that the metabolic modelling was a good predictor of biomass in the context of maize-*R. irregularis* interaction especially for B73 maize genotype. The MBA analysis was able to predict that pyrimidine biosynthesis is crucial in this interaction, in agreement with experimental observation of pyrimidine accumulation in leaves of symbiotic Lo3 plants under LN.

**Table 1:**
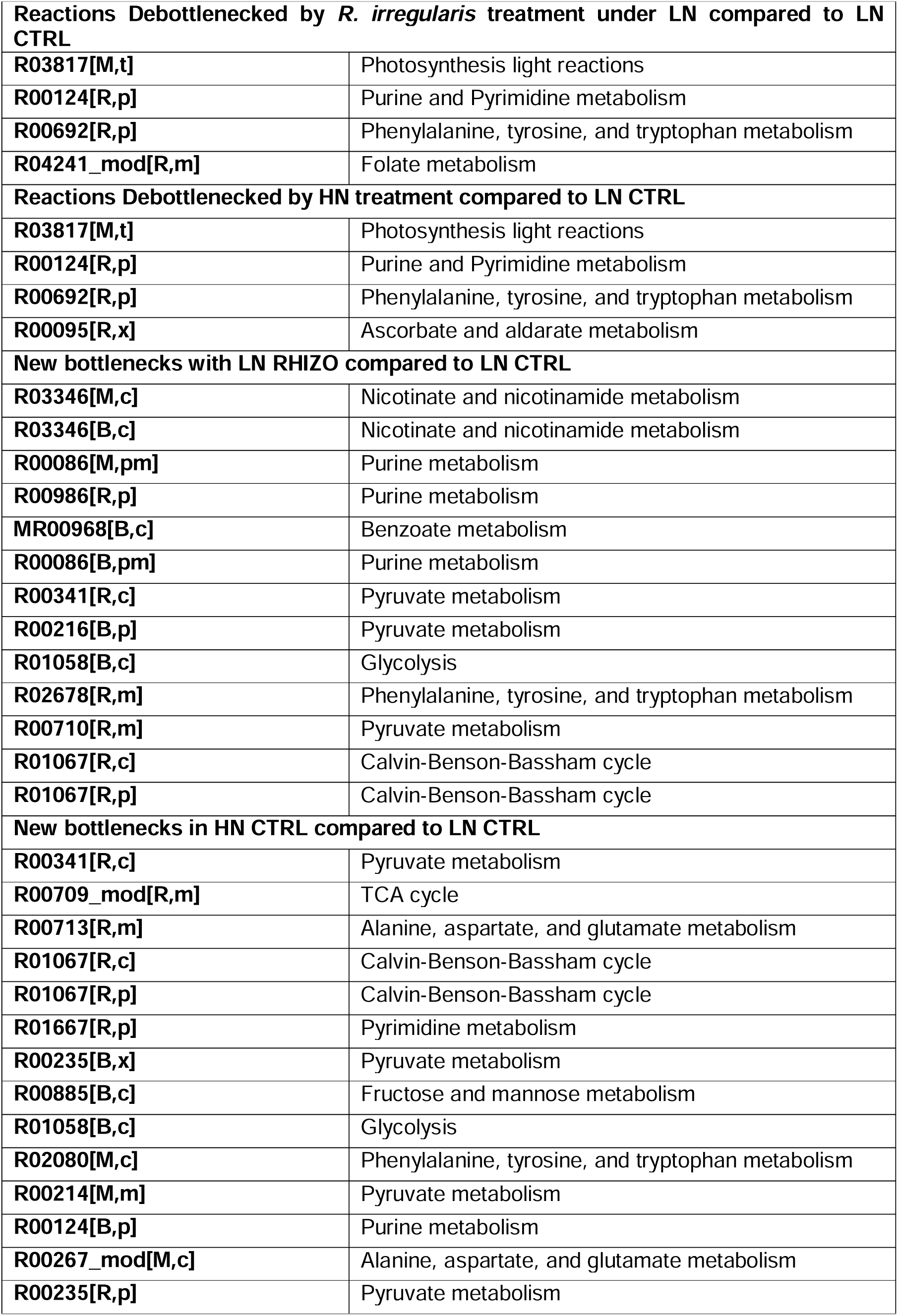
List of metabolic bottlenecks relieved and created by *R. irregularis* or N supply in B73. Metabolic bottlenecks were identified by MBA that are relieved by either R. irregularis treatment or HN treatment compared to LN control conditions are indicated as “debottlnecked”. Those that were newly created following R. irregularis or HN treatment were compared to LN control conditions are indicated as “new metabolic” bottelnecks. M: mesophyll, R: root, p: plastid, t: thylakoid, c: cytosol, x: vacuole.

| Reactions Debottlenecked by *R. irregularis* treatment under LN compared to LN CTRL |  |
| --- | --- |
| R03817[M,t] | Photosynthesis light reactions |
| R00124[R,p] | Purine and Pyrimidine metabolism |
| R00692[R,p] | Phenylalanine, tyrosine, and tryptophan metabolism |
| R04241_mod[R,m] | Folate metabolism |
| Reactions Debottlenecked by HN treatment compared to LN CTRL |  |
| R03817[M,t] | Photosynthesis light reactions |
| R00124[R,p] | Purine and Pyrimidine metabolism |
| R00692[R,p] | Phenylalanine, tyrosine, and tryptophan metabolism |
| R00095[R,x] | Ascorbate and aldarate metabolism |
| New bottlenecks with LN RHIZO compared to LN CTRL |  |
| R03346[M,c] | Nicotinate and nicotinamide metabolism |
| R03346[B,c] | Nicotinate and nicotinamide metabolism |
| R00086[M,pm] | Purine metabolism |
| R00986[R,p] | Purine metabolism |
| MR00968[B,c] | Benzoate metabolism |
| R00086[B,pm] | Purine metabolism |
| R00341[R,c] | Pyruvate metabolism |
| R00216[B,p] | Pyruvate metabolism |
| R01058[B,c] | Glycolysis |
| R02678[R,m] | Phenylalanine, tyrosine, and tryptophan metabolism |
| R00710[R,m] | Pyruvate metabolism |
| R01067[R,c] | Calvin-Benson-Bassham cycle |
| R01067[R,p] | Calvin-Benson-Bassham cycle |
| New bottlenecks in HN CTRL compared to LN CTRL |  |
| R00341[R,c] | Pyruvate metabolism |
| R00709_mod[R,m] | TCA cycle |
| R00713[R,m] | Alanine, aspartate, and glutamate metabolism |
| R01067[R,c] | Calvin-Benson-Bassham cycle |
| R01067[R,p] | Calvin-Benson-Bassham cycle |
| R01667[R,p] | Pyrimidine metabolism |
| R00235[B,x] | Pyruvate metabolism |
| R00885[B,c] | Fructose and mannose metabolism |
| R01058[B,c] | Glycolysis |
| R02080[M,c] | Phenylalanine, tyrosine, and tryptophan metabolism |
| R00214[M,m] | Pyruvate metabolism |
| R00124[B,p] | Purine metabolism |
| R00267_mod[M,c] | Alanine, aspartate, and glutamate metabolism |
| R00235[R,p] | Pyruvate metabolism |

**Table 2:** List of metabolic bottlenecks relieved and created by *R. irregularis* or N supply on Lo3. Metabolic bottlenecks were identified by MBA that are relieved by either *R. irregularis* treatment or HN treatment compared to LN control conditions are indicated as “debottlenecked”. Those that were newly created following *R. irregularis* or HN treatment were compared to LN control conditions are indicated as “new metabolic” bottlenecks. M: mesophyll, R: root, p: plastid, t: thylakoid, c: cytosol, x: vacuole.

| **Reactions Debottlenecked by *R. irregularis* treatment under LN compared to LN CTRL** |  |
| --- | --- |
| **R01080[R,p]** | Pyrimidine metabolism |
| **R02143[R,p]** | Purine metabolism |
| **R01667[R,p]** | Pyrimidine metabolism |
| **R01067[R,c]** | Calvin-Benson-Bassham cycle |
| **R01067[R,p]** | Calvin-Benson-Bassham cycle |
| **R00281[R,p]** | Non pathway information |
| **Reactions Debottlenecked by HN treatment compared to LN CTRL** |  |
| **R04241_mod[R,m]** | Folate metabolism |
| **R05395[R,p]** | DDT degradation |
| **New bottlenecks with LN RHIZO compared to LN CTRL** |  |
| **R00709_mod[R,m]** | TCA cycle |
| **R00713[R,m]** | Alanine, aspartate, and glutamate metabolism |
| **MR00886[R,m]** | Acyl carrier protein metabolism |
| **New bottlenecks in HN CTRL compared to LN CTRL** |  |
| **R00709_mod[R,m]** | TCA cycle |
| **R00713[R,m]** | Alanine, aspartate, and glutamate metabolism |
| **R00885[B,c]** | Fructose and mannose metabolism |

### Fungal transcriptome of in inoculated maize roots is driven by N availability

We then investigated the metabolic cross talk between *R. irregularis* and maize from th fungal side. For this purpose, we determined the fungal genes displaying a higher expression in LN roots compared to HN roots and vice versa. This selection was performed for B73 and Lo3 separately. The Venn diagrams in Figure 6A and 6B show the numbers of fungal genes up-regulated under LN compared to HN or vice versa in each maize line. In both maize lines, the functional categories overrepresented in *R. irregularis* DEG were similar in each N nutrition condition Figure 6C. Among the DEG functional categories, several genes were involved in amino acid and nucleotide metabolism and transport as well as in inorganic transport. We focused on the expression profiles of nucleotide and amino acid metabolism genes and transporter genes as well as phosphate and ammonium acid transport genes in the annotation database of JGI (https://genome.jgi.doe.gov/portal/). Supplementary Table 6 shows the expression pattern of these fungal genes. Interestingly, fungal ammonia transporters are more highly expressed under LN than under HN. The fungal amino acid biosynthetic genes up-regulated are mainly involved in asparagine, arginine, glutamine, glutamate and serine pathways. Several fungal amino acid transporter genes were found to be up-regulated either in LN or HN roots. The fungal genes related to pyrimidine de novo biosynthesis and ureide metabolism (Figure 6D) are expressed in both LN and HN roots. Interestingly, although expressed under all nutritional conditions they are globally up-regulated in HN roots. These data show that the fungal transcriptome is strongly affected by the nutritional N status of the host and the soil and that fungal amino acid and pyrimidine metabolism genes are expressed in symbiotic roots.

**Figure 6:**
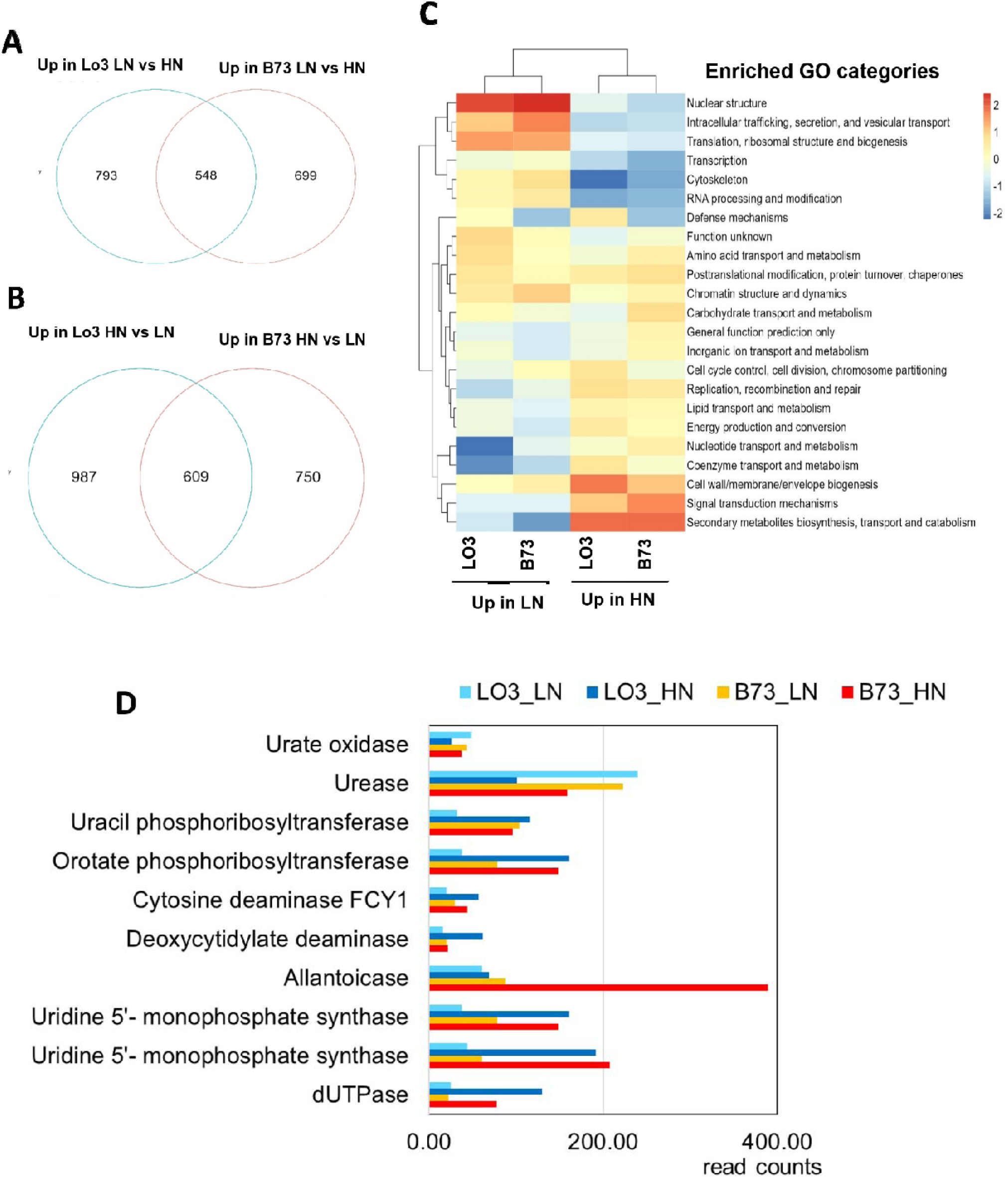
N supply strongly impacts *Rhlzophegus lrregularls* transcriptome and pyrimidine biosynthetic genes in maize roots. Fungal DEGs were aalectad so that they have a fold change (FC >1.5). **A:** Up in LN Indicates ***R.*** *lrreguJarls* genes that are more highly expressed In Inoculated LN roots compared to Inoculated HN roots. **B:** Up In HN Indicates *R. lrregu/erls* genes that are more highly expreasecl In HN compared to LN, C: Gene ontology of functional categories enrlchmenl was based on KOO annotations of encoded *R. lrregularls* proteins (see materials and methods), Each bar represents the fUnctlonal category enriched In the given condition. p-value of enrichment was < 0.05; only categories enriched more than 2 fold and dlsplaylng more lhan 2 genes are represented and used to build the haatmap. The color scale Indicates the over-representaUon or the down-representation In red and blue respectively. **D:** read counts of *R. lm,guJarts* transcripts related to pyrimidine de novo blosynthesis and salvage (from supplementary Table 6).

## Discussion

In our study, we showed that a symbiotic association with *R. irregularis* was able to compensate for decreased maize plant performance when N fertilization is reduced. In the literature, depending on both the N supply level and the form of N supplied, different effects on yield and/or biomass production have been reported, including an increase, no effect, or a reduction in productivity when the plant develops a symbiotic association with an AMF ^26,58,59^. In the present investigation, we estimated that kernel yield can be maintained with a 2– to 5-fold reduction in N input when two maize lines were inoculated with *R. irregularis*. Such a beneficial impact was previously observed for maize plant biomass production under low N fertilizer inputs ^26^. One can, therefore, conclude that the positive impact of AMF on plant performance can be variable depending on the AMF and the host genotype as well as the level and type of available N source ^9,13,58^.

Our transcriptome study allowed us to provide an overview of biological processes putatively involved during the establishment of a symbiotic association between the AMF *R. irregularis* and maize. The DEGs that were either up or down-regulated in roots were enriched in transcripts encoding proteins involved in a variety of biological functions: defence, hormone signalling N and C metabolism, cell wall and chromatin remodelling, proteolysis, metal homeostasis and lipid metabolism. Similar results were previously obtained in several other model and crop species ^27,60–67^. In particular, the most highly up-regulated genes in roots of B73 and Lo3 under both N nutrition conditions in response to *R. irregularis* inoculation are encoding peptidases, peptidase inhibitors, chitinases and senescence genes (Supplementary Table 7a and b). Interestingly, autophagy genes are overrepresented in Lo3 genes up-regulated in roots under LN following inoculation with *R. irregularis* (Supplementary Table 8, Supplementary Figure 7). These data suggest that important proteolysis processes are taking place in in inoculated roots that may involve autophagy especially in the Lo3 maize line. Autophagy is a process known to recycle N during senescence, and is known to provide N assimilates necessary for maize kernel filling ^68,69^.

Changes in the level of expression of genes involved in sugar, amino acid, and lipid metabolism were confirmed by metabolomics analyses (Supplementary Figure 9 and 10). Globally, we observed lipid accumulation in roots colonized by AMFs and a drop in soluble sugars in both roots and shoots, a phenomenon already described ^16,70^. This is considered a way of fuelling the symbiotic fungus with a carbon source as lipids. In inoculated plants, arginine (Arg), known to be a key molecule involved in the transfer of N from the fungus to the plant ^11,13,66,71^ accumulates under HN conditions not only in the roots of line B73 HN but also in the leaves.

To delve deeper than a mere data overview, the incorporation of transcriptomics data into the iZMA6517 metabolic model is especially potent, as we observed concurrence between iZMA6517 predictions and actual experimental results regarding biomass increase triggered by the AMF. Our previous efforts of maize metabolic modelling resulted in excellent phenotypical predictions for including the photosynthetic behaviour of maize leaf ^32^, predicting the root phenotype under nitrogen-optimal and nitrogen deficient conditions^33^ and whole plant phenotype for heat and cold stress conditions ^34^. Thereby, our whole-plant model of maize, iZMA6517 ^34^ was an ideal candidate to identify metabolic underpinnings of maize for different nitrogen availability. Interestingly, iZMA6517 allowed the identification of pyrimidine metabolism as a crucial player in this symbiosis. Indeed, our transcriptomic analysis demonstrates that, regardless of nitrogen feeding conditions, pyrimidine biosynthesis processes are consistently downregulated in leaves of inoculated plants (Supplementary Figure 7, Supplementary Table 10). Under LN conditions, several key genes encoding enzymes involved in nucleotide metabolism, including putative Orotidine-5’-phosphate decarboxylase, thiaminase, Uridine 5’-monophosphate synthase, Cytidine triphosphate synthase, are downregulated in leaves (Supplementary Table 10). A critical gene in this context is *Zm00001d023833*, encoding a putative nucleoside diphosphate kinase (NDPK), which participates in pyrimidine and purine nucleotide salvage ^72^. This enzyme was identified as a metabolic bottleneck relieved by AMF inoculation (Figure 5, Table 1 and 2) and its expression in the context of symbiosis was predicted to lead to accumulation of pyrimidine in leaves. Remarkably, this accumulation was confirmed experimentally by the accumulation of UMP and UDP in Lo3 leaves of LN symbiotic plants (Figure 5D).

In line with these data, fungal transcriptomic analysis by RNAseq shows that pyrimidine and ureide metabolism are expressed by *R. irregularis*. The integration of maize and fungal RNA-seq and metabolome analyses has allowed us to construct a simplified model depicting the various metabolic N related events occurring in the symbiotic association considering both lines (Figure 7, Supplementary Figure 11). Under LN, fungal argininosuccinate lyase, carbamoyl-phosphate synthase and arginase genes are upregulated, potentially leading to urea accumulation. However, urea level drops in LN inoculated roots (Supplementary Figure 11), a fact that is consistent with higher expression of *R. irregularis* urease under LN (Figure 6, Supplementary Table 6). In parallel, fungal genes involved in pyrimidine de novo synthesis and salvage are expressed in both Lo3 and B73 symbiotic roots under LN and HN (Figure 6, Supplementary Table 6). Our hypothesis is that pyrimidines are transferred to the leaves via the phloem sap, and that they are provided by the fungus especially under LN. This is consistent with the finding that in Lo3 leaves of symbiotic plants, UMP, UDP and UDP-glc levels were higher compared to control plants. The fact that in B73 leaves, and in HN leaves, we did not observe an increase of these metabolites in response to AMF inoculation indicates that their accumulation is tightly controlled. In our experiment the level of pyrimidines was globally higher in HN leaves than LN leaves but no impact of AMF inoculation was observed in HN plants (Supplementary Figure 12) indicating that the steady state of these compounds may vary over time. On the other hand, allantoin accumulates exclusively in HN roots and its level remains unaffected by inoculation (Supplementary Figure 11). This metabolite can be converted into allantoic acid, which may be released into the apoplast ^73^. A fungal allantoicase is upregulated in HN roots, indicating possible conversion of allantoic acid into urea under HN conditions. This transfer of ureides is akin to the process observed in symbiotic associations between tropical legumes and Rhizobia species ^73^. Previous studies have also identified upregulation of genes involved in allantoin synthesis in roots of AMF-inoculated plants ^74^. Consequently, it is plausible that ureide transfer mirrors that occurring in legume nodules.

**Figure 7:**
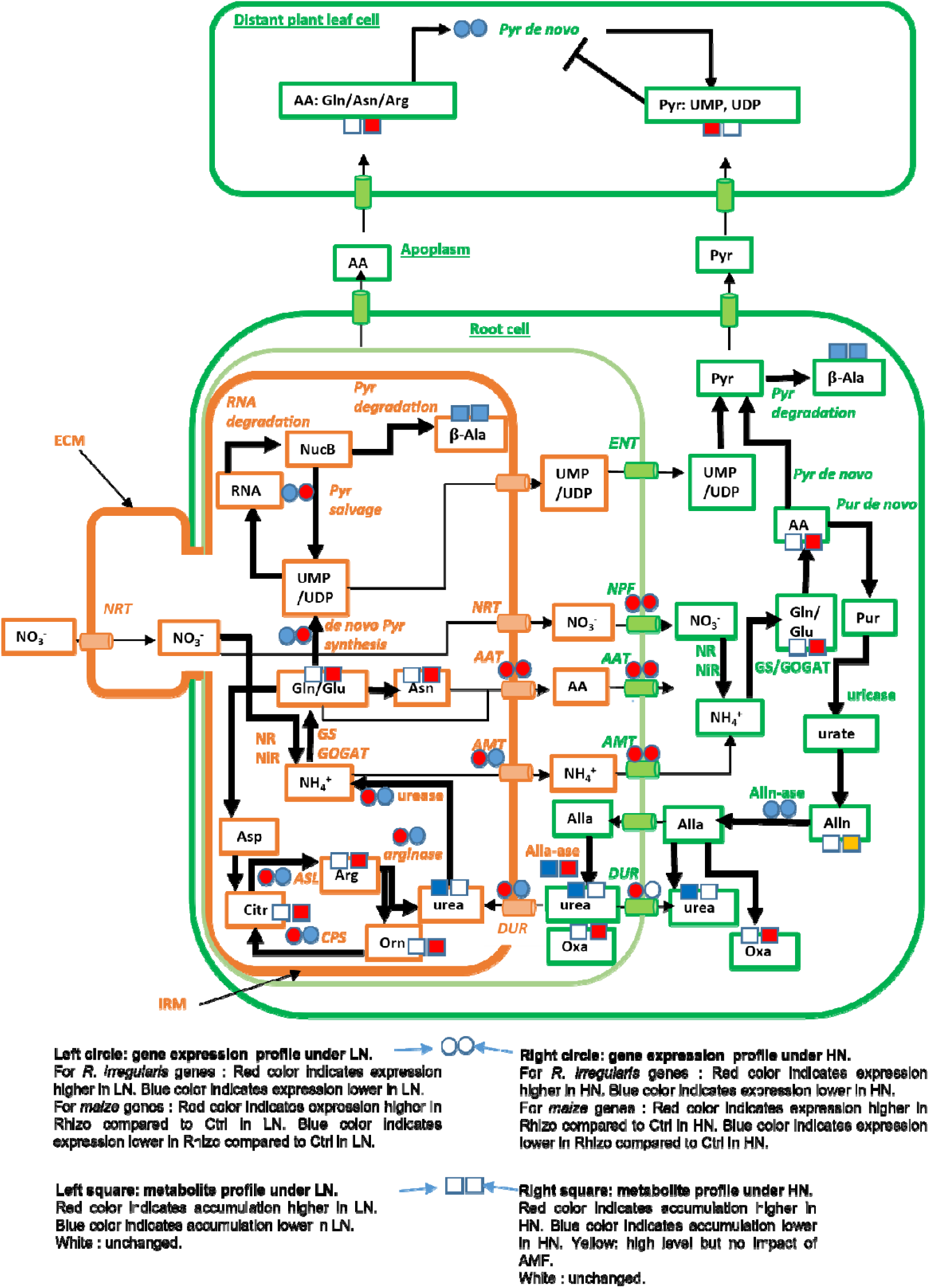
H**y**pothetical **working model for maize and *R. irregularis* N nutrition under LN and HN.** To build the model, the metabolomic data from B73 and Lo3 maize lines were pooled for simplification (Supplementary Figure 10) and we used genes in Supplementary Tables 6 and 8. In some cases genes with multigene families such as Glutamine Synthetase encoding genes have different expression profiles. For simplification purposes, these cases were not considered in the hypothetical mechanistic model below. Genes are indicated in italics. Metabolites are in boxes. Surrounded by orange boxes: fungal metabolites. Surrounded by green boxes: plant metabolites. Written in green or orange are plant or fungal genes respectively. If no box or circle is indicated, this means that there are no clear data bout the corresponding gene or metabolite. AA: Amino acids. NucB: Nucleobases. Pur: Purine. Pyr: Pyrimidine. IRM: Intraradical mycelium. ECM: Extraradical mycelium. Alla: Allantoic Acid. Alln: Allantoin. Alln-ase: Allantoinase. Arg: arginine. Asn: Asparagine. Asp: Aspartate. b-Ala:b-alanine. Citr: Citrulline. Glu: Glutamate. Gln: Glutamine. Orn: Ornithine. GOGAT: Glutamine Oxo-Glutarate Aminotransferase. Oxa: Oxalic Acid. NR: Nitrate reductase. NiR: Nitrite reductase. AAT: Amino acid transporter. AMT: Ammonium transporter. DUR: Urea transporter. NPF: plant Nitrate transporter. NRT: *R. irregularis* nitrate transporter. CPS: Carbamoyl-phosphate synthase. ASL: Argininosuccinate lyase. Pyr de novo: Pyrimidine de novo biosynthetic pathway. Pyr degradation: Pyrimidine degradation. Pur de novo: Pur de novo biosynthetic pathway.

Fungal arbuscules are known to undergo controlled senescence ^17,22^. Our data suggest that autophagy and proteolysis in general are upregulated in mycorrhizal maize roots, with higher expression in Lo3 LN roots (Figure 3 and Supplementary Table 8). Autophagy may contributes to the degradation of fungal cellular products, potentially releasing nucleosides. The fungal nucleotide salvage process can then recycle nucleosides that are probably transported to the leaves thus repressing de novo plant pyrimidine biosynthesis pathway (Supplementary Figure 7, Supplementary Table 9) ^72,73^. This implies that AMF likely supplies the host plant with nucleotides and nucleosides, potentially pyrimidines, to circumvent the deficiency in NDPK activity, thereby alleviating the associated metabolic bottleneck. This observation is consistent with the known role of NDPK in plant survival with respect to pyrimidine salvage ^72,73^. It is noteworthy that pyrimidine biosynthesis can be an essential process for plant N management ^75,76^. Pyrimidine metabolism holds significance not only in crop performance but also in nitrogen use efficiency (NUE) ^73,77,78^. Recent findings demonstrate a correlation between maize leaf nitrogen accumulation and NDPK protein levels ^79^, further supporting the hypothesis that *R. irregularis* stimulates the reprogramming of plant pyrimidine metabolism to enhance maize performance under nitrogen-limiting conditions.

Physiological profiling combined with metabolic modelling revealed pyrimidine and ureide metabolism as previously unrecognized pivotal components in this interaction, warranting additional investigation ^80,81^. Additionally, the contrasting strategies adopted by our two maize lines to capitalize on the fungus highlight the significance of harnessing maize’s broad genetic diversity for a holistic comprehension of this symbiotic relationship ^24,82,83^.

## Data Availability

RNAseq project is deposited (Gene Expression Omnibus,Edgard R. et al. 2002): https://www.ncbi.nlm.nih.gov/geo/query/acc.cgi?acc=GSE235654; accession no. GSE235654. All steps of the experiment, from growth conditions to bioinformatic analyses, were detailed in CATdb ^84^: http://tools.ips2.u-psud.fr.fr/CATdb/; Project: NGS2021_19_Rhizophagus according to the MINSEQE ‘minimum information about a high throughput sequencing experiment’.

## Acknowledgments

BD acknowledges Université Paris Saclay for PhD fellowship. AD acknowledges the benefits received from France 2030 program (Saclay Plant Sciences, reference n° ANR-17-EUR-0007, EUR SPS-GSR integrated into France 2030 (reference n° ANR-11-IDEX-0003-02), INRAE Grant IB BAP 2021 MYCORN, and IJPB’s Plant Observatory technological platforms. RS gratefully acknowledges funding support from National Science Foundation (NSF) CAREER grant (1943310) and Nebraska Collaboration Initiative Grant (21-1106-6011). We thank Michel Lebrusq, Lilan Diahuron, Patrick Grillot, and Christian Jeudy for taking care of the plants in the greenhouse. We thank the POPS platform for the RNAseq experiment and data. We also thank Etienne Delannoy from POPS platform for fruitful discussions. We are grateful to Cyril Bauland and Carine Palaffre from the INRAE for providing seeds from the Centre de Ressources Biologiques of Saint Martin de Hinx (France). We acknowledge Pr. Mario MOTTO from ISC, Bergamo (Italy) for providing Lo3 seeds.

## Notes

### Competing Interest Statement

The authors have declared no competing interest.

### Summary of Updates

This version has been revised essentially to shorten the text and the Figures as well as to include new data sets for metabolic modeling for the Lo3 line and monitoring of pyrimidine content that confirms the metabolic bottleneck analyses.

