## Supplementary Figure 1 for "Unlocking the Mycorrhizal Nitrogen Pathway Puzzle: Metabolic Modelling and multi-omics unveil Pyrimidines’ Role in Maize Nutrition via Arbuscular Mycorrhizal Fungi Amidst Nitrogen Scarcity"

### Slide 1
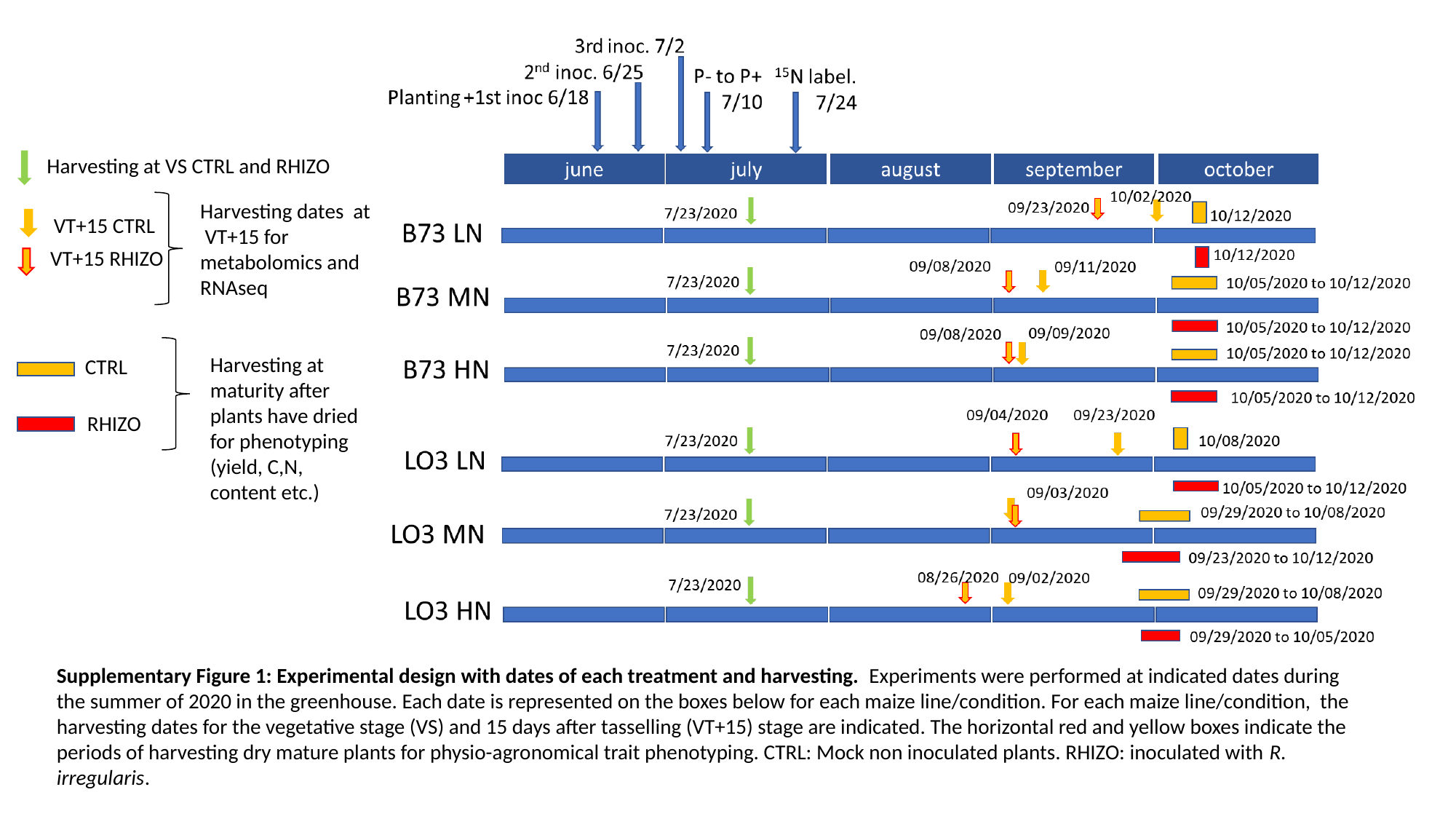

Harvesting at VS CTRL and RHIZO
Harvesting dates at VT+15 for metabolomics and
RNAseq
VT+15 CTRL
VT+15 RHIZO
Harvesting at maturity after plants have dried for phenotyping (yield, C,N, content etc.)
CTRL
RHIZO
Supplementary Figure 1: Experimental design with dates of each treatment and harvesting. Experiments were performed at indicated dates during the summer of 2020 in the greenhouse. Each date is represented on the boxes below for each maize line/condition. For each maize line/condition, the harvesting dates for the vegetative stage (VS) and 15 days after tasselling (VT+15) stage are indicated. The horizontal red and yellow boxes indicate the periods of harvesting dry mature plants for physio-agronomical trait phenotyping. CTRL: Mock non inoculated plants. RHIZO: inoculated with R. irregularis.
