## Supplementary Figure 3 for "Unlocking the Mycorrhizal Nitrogen Pathway Puzzle: Metabolic Modelling and multi-omics unveil Pyrimidines’ Role in Maize Nutrition via Arbuscular Mycorrhizal Fungi Amidst Nitrogen Scarcity"

### Slide 1
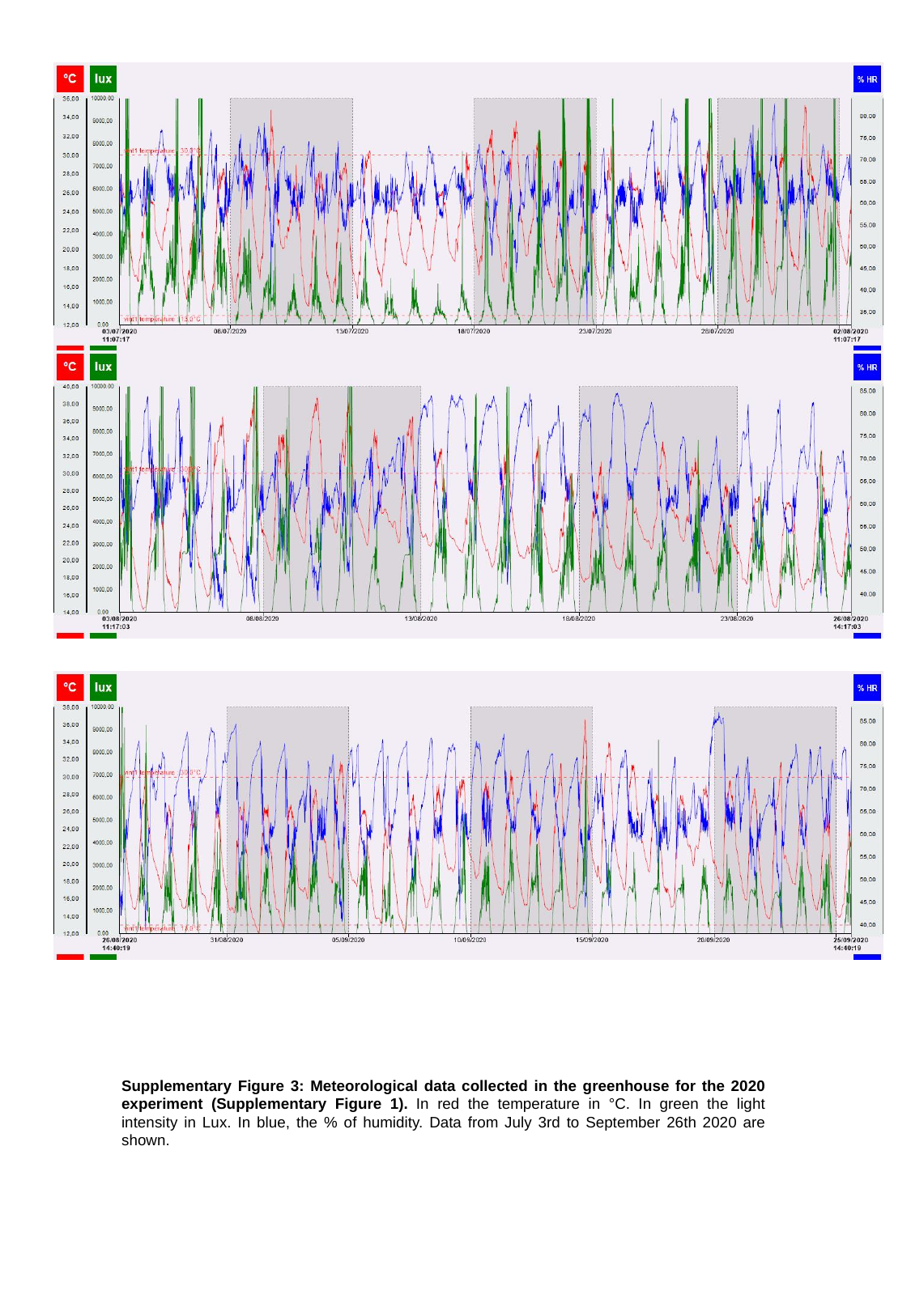

Supplementary Figure 3: Meteorological data collected in the greenhouse for the 2020 experiment (Supplementary Figure 1). In red the temperature in °C. In green the light intensity in Lux. In blue, the % of humidity. Data from July 3rd to September 26th 2020 are shown.
