## Supplementary Figure 4 for "Unlocking the Mycorrhizal Nitrogen Pathway Puzzle: Metabolic Modelling and multi-omics unveil Pyrimidines’ Role in Maize Nutrition via Arbuscular Mycorrhizal Fungi Amidst Nitrogen Scarcity"

### Slide 1
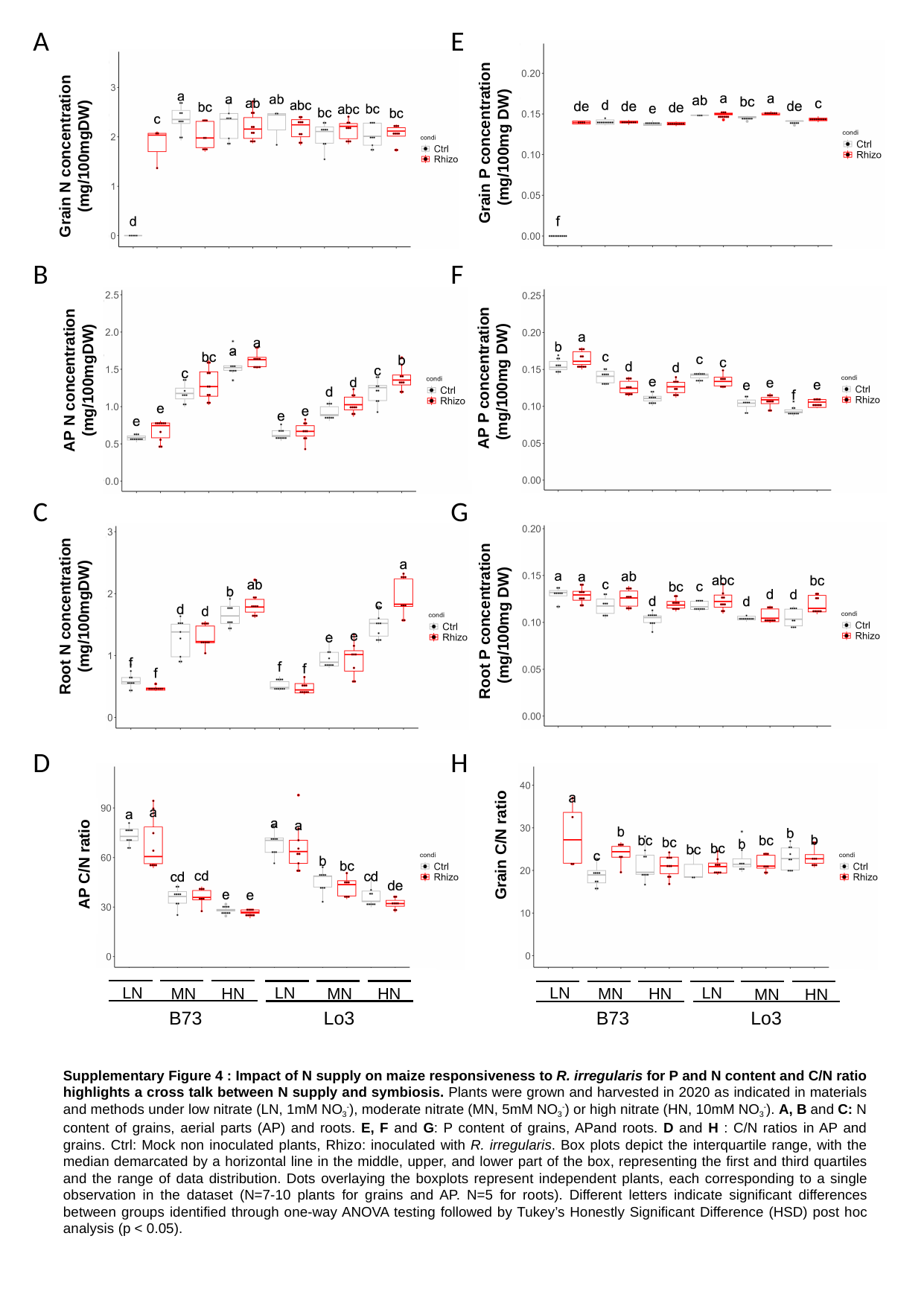

A
E
 Grain P concentration
(mg/100mg DW)
 Grain N concentration
(mg/100mgDW)
B
F
 AP P concentration
(mg/100mg DW)
 AP N concentration
(mg/100mgDW)
C
G
 Root N concentration
(mg/100mgDW)
 Root P concentration
(mg/100mg DW)
D
H
 Grain C/N ratio
 AP C/N ratio
LN
MN
HN
B73
LN
MN
HN
Lo3
LN
MN
HN
B73
LN
MN
HN
Lo3
Supplementary Figure 4 : Impact of N supply on maize responsiveness to R. irregularis for P and N content and C/N ratio highlights a cross talk between N supply and symbiosis. Plants were grown and harvested in 2020 as indicated in materials and methods under low nitrate (LN, 1mM NO_3_^-^), moderate nitrate (MN, 5mM NO_3_^-^) or high nitrate (HN, 10mM NO_3_^-^). A, B and C: N content of grains, aerial parts (AP) and roots. E, F and G: P content of grains, APand roots. D and H : C/N ratios in AP and grains. Ctrl: Mock non inoculated plants, Rhizo: inoculated with R. irregularis. Box plots depict the interquartile range, with the median demarcated by a horizontal line in the middle, upper, and lower part of the box, representing the first and third quartiles and the range of data distribution. Dots overlaying the boxplots represent independent plants, each corresponding to a single observation in the dataset (N=7-10 plants for grains and AP. N=5 for roots). Different letters indicate significant differences between groups identified through one-way ANOVA testing followed by Tukey’s Honestly Significant Difference (HSD) post hoc analysis (p < 0.05).
