## Supplementary Figure 5 for "Unlocking the Mycorrhizal Nitrogen Pathway Puzzle: Metabolic Modelling and multi-omics unveil Pyrimidines’ Role in Maize Nutrition via Arbuscular Mycorrhizal Fungi Amidst Nitrogen Scarcity"

### Slide 1
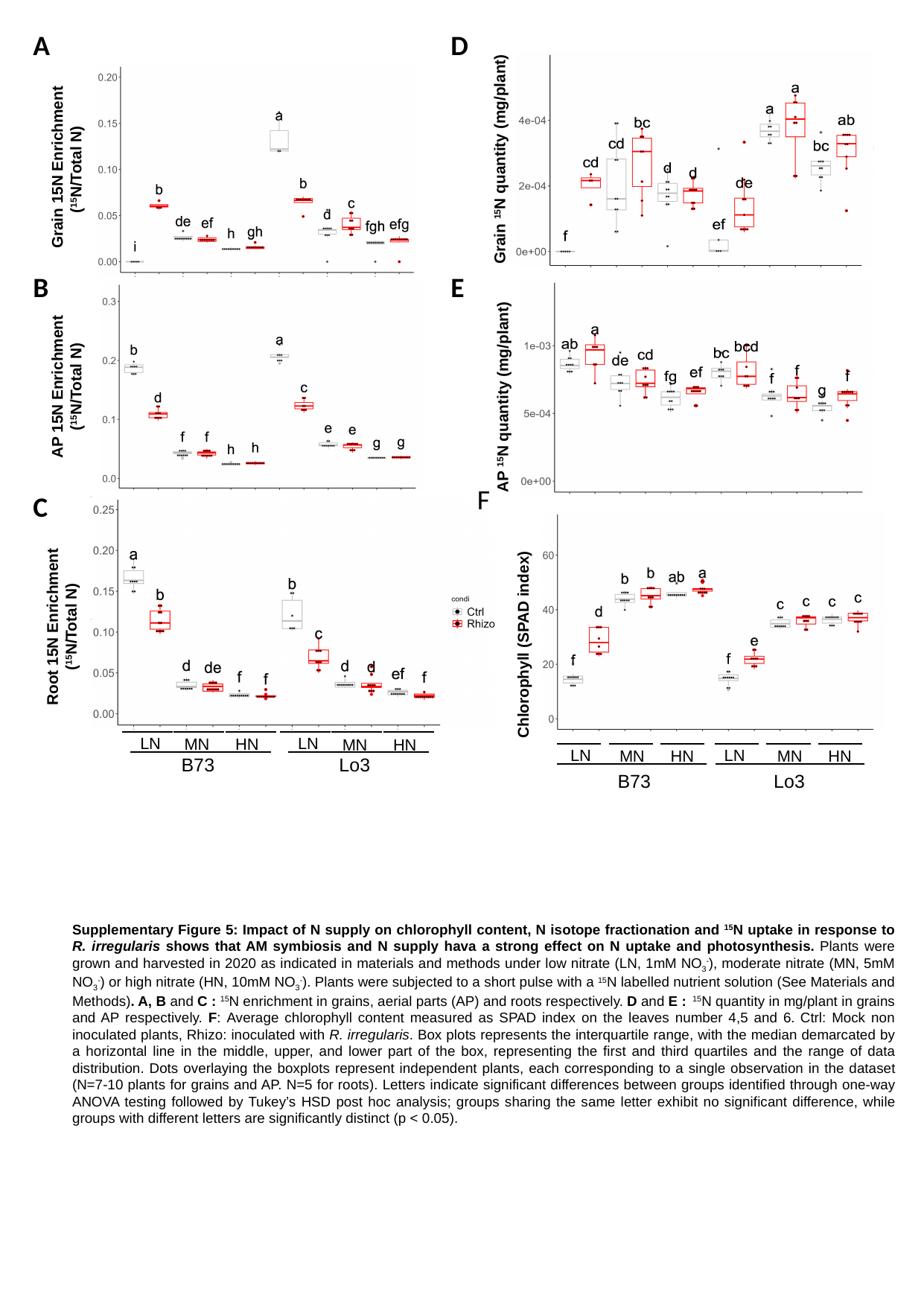

A
D
Grain ^15^N quantity (mg/plant)
 Grain 15N Enrichment (^15^N/Total N)
B
E
E
AP 15N Enrichment (^15^N/Total N)
AP ^15^N quantity (mg/plant)
F
C
Root 15N Enrichment (^15^N/Total N)
Chlorophyll (SPAD index)
Ctrl
Rhizo
Ctrl
Rhizo
Ctrl
LN
MN
HN
B73
Rhizo
Ctrl
Rhizo
Ctrl
Rhizo
Ctrl
LN
MN
HN
Lo3
Rhizo
LN
MN
HN
B73
LN
MN
HN
Lo3
Supplementary Figure 5: Impact of N supply on chlorophyll content, N isotope fractionation and ^15^N uptake in response to R. irregularis shows that AM symbiosis and N supply hava a strong effect on N uptake and photosynthesis. Plants were grown and harvested in 2020 as indicated in materials and methods under low nitrate (LN, 1mM NO_3_^-^), moderate nitrate (MN, 5mM NO_3_^-^) or high nitrate (HN, 10mM NO_3_^-^). Plants were subjected to a short pulse with a ^15^N labelled nutrient solution (See Materials and Methods). A, B and C : ^15^N enrichment in grains, aerial parts (AP) and roots respectively. D and E : ^15^N quantity in mg/plant in grains and AP respectively. F: Average chlorophyll content measured as SPAD index on the leaves number 4,5 and 6. Ctrl: Mock non inoculated plants, Rhizo: inoculated with R. irregularis. Box plots represents the interquartile range, with the median demarcated by a horizontal line in the middle, upper, and lower part of the box, representing the first and third quartiles and the range of data distribution. Dots overlaying the boxplots represent independent plants, each corresponding to a single observation in the dataset (N=7-10 plants for grains and AP. N=5 for roots). Letters indicate significant differences between groups identified through one-way ANOVA testing followed by Tukey’s HSD post hoc analysis; groups sharing the same letter exhibit no significant difference, while groups with different letters are significantly distinct (p < 0.05).
