## Supplementary Figure 6 for "Unlocking the Mycorrhizal Nitrogen Pathway Puzzle: Metabolic Modelling and multi-omics unveil Pyrimidines’ Role in Maize Nutrition via Arbuscular Mycorrhizal Fungi Amidst Nitrogen Scarcity"

### Slide 1
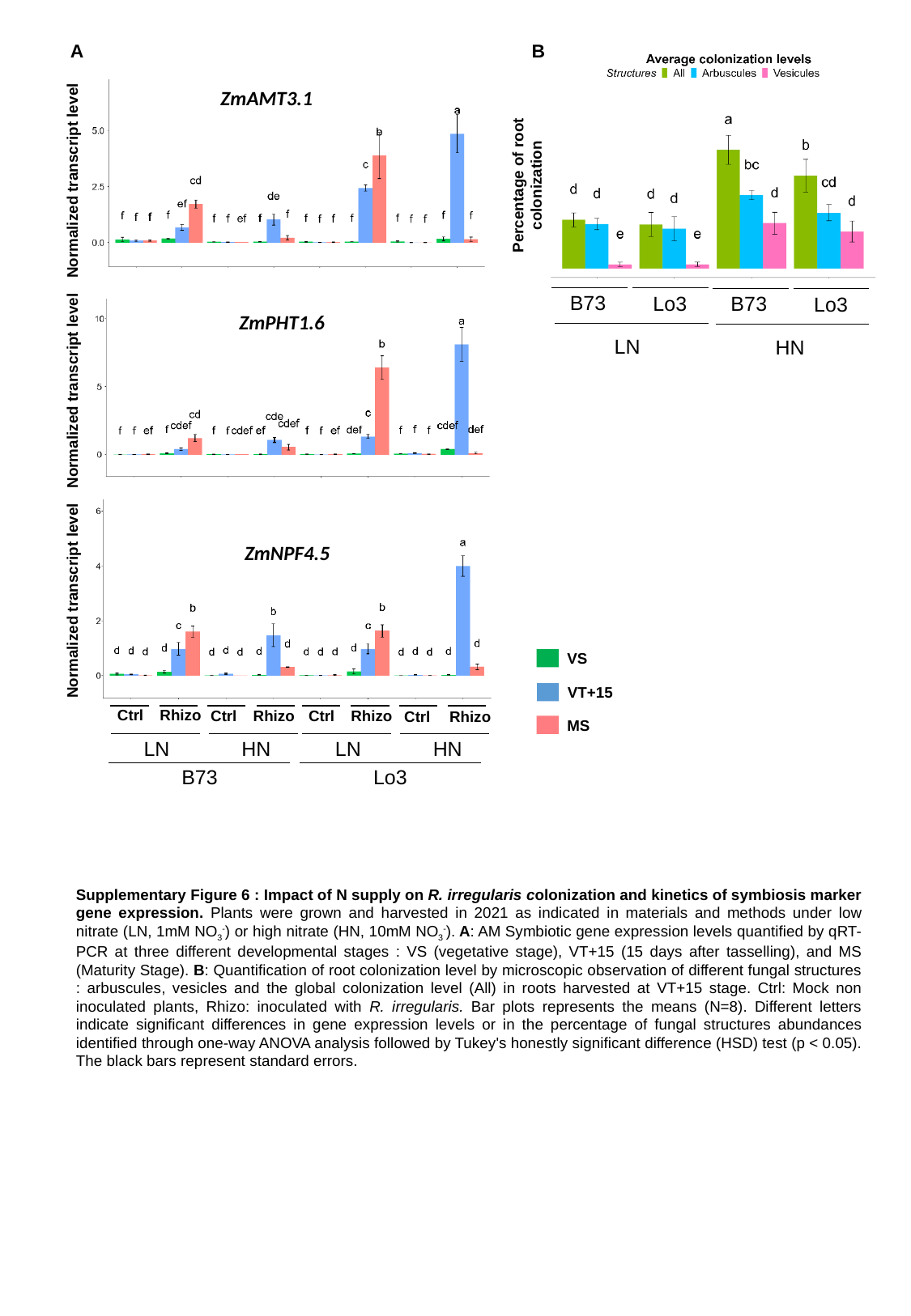

A
B
Percentage of root colonization
B73
B73
Lo3
Lo3
LN
HN
ZmAMT3.1
Normalized transcript level
ZmPHT1.6
Normalized transcript level
ZmNPF4.5
Normalized transcript level
VS
VT+15
Ctrl
Rhizo
Ctrl
Rhizo
Ctrl
Rhizo
Ctrl
Rhizo
MS
LN
HN
LN
HN
B73
Lo3
Supplementary Figure 6 : Impact of N supply on R. irregularis colonization and kinetics of symbiosis marker gene expression. Plants were grown and harvested in 2021 as indicated in materials and methods under low nitrate (LN, 1mM NO_3_^-^) or high nitrate (HN, 10mM NO_3_^-^). A: AM Symbiotic gene expression levels quantified by qRT-PCR at three different developmental stages : VS (vegetative stage), VT+15 (15 days after tasselling), and MS (Maturity Stage). B: Quantification of root colonization level by microscopic observation of different fungal structures : arbuscules, vesicles and the global colonization level (All) in roots harvested at VT+15 stage. Ctrl: Mock non inoculated plants, Rhizo: inoculated with R. irregularis. Bar plots represents the means (N=8). Different letters indicate significant differences in gene expression levels or in the percentage of fungal structures abundances identified through one-way ANOVA analysis followed by Tukey's honestly significant difference (HSD) test (p < 0.05). The black bars represent standard errors.
