## Supplementary Figure 7 for "Unlocking the Mycorrhizal Nitrogen Pathway Puzzle: Metabolic Modelling and multi-omics unveil Pyrimidines’ Role in Maize Nutrition via Arbuscular Mycorrhizal Fungi Amidst Nitrogen Scarcity"

### Slide 1
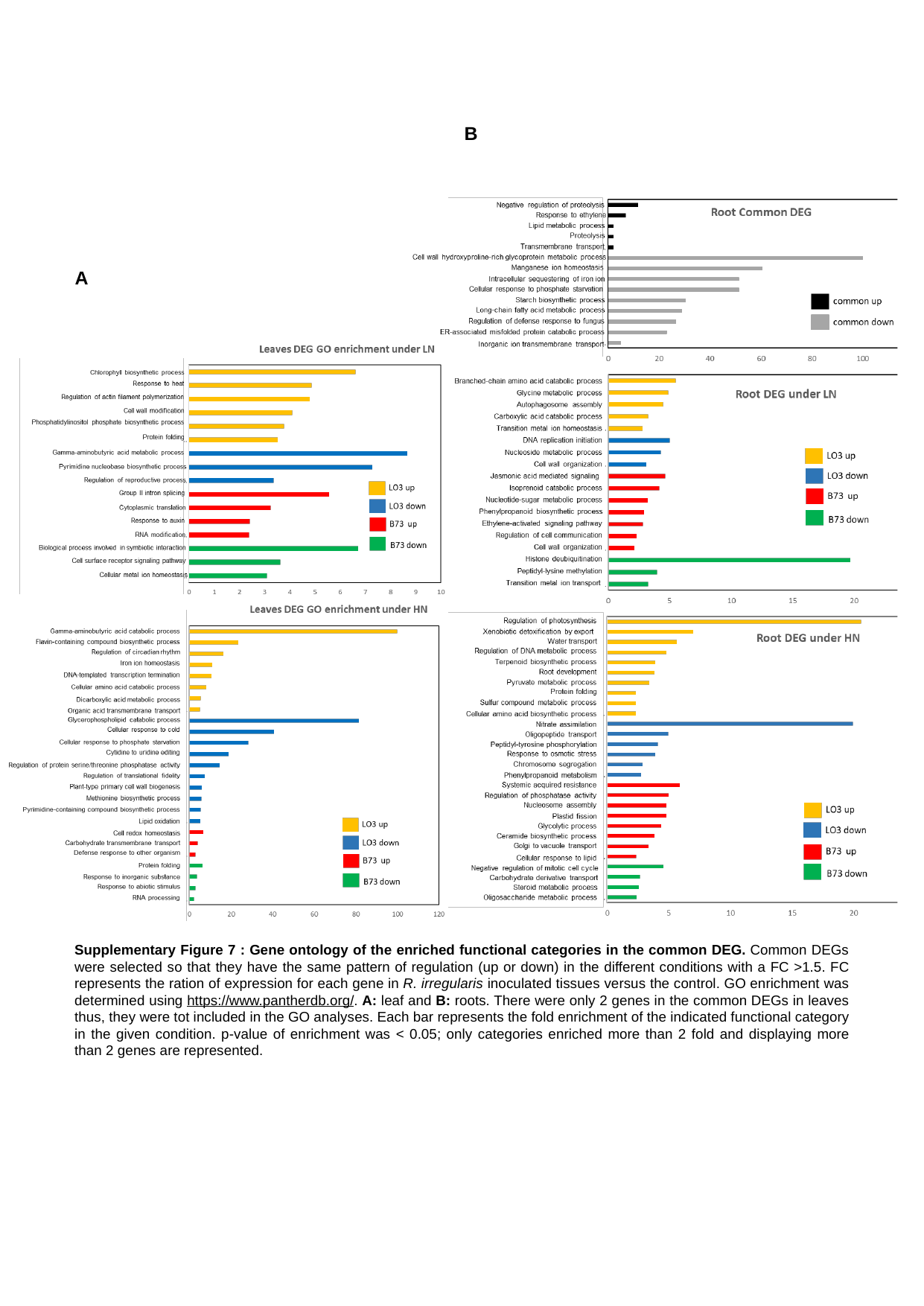

B
A
Supplementary Figure 7 : Gene ontology of the enriched functional categories in the common DEG. Common DEGs were selected so that they have the same pattern of regulation (up or down) in the different conditions with a FC >1.5. FC represents the ration of expression for each gene in R. irregularis inoculated tissues versus the control. GO enrichment was determined using https://www.pantherdb.org/. A: leaf and B: roots. There were only 2 genes in the common DEGs in leaves thus, they were tot included in the GO analyses. Each bar represents the fold enrichment of the indicated functional category in the given condition. p-value of enrichment was < 0.05; only categories enriched more than 2 fold and displaying more than 2 genes are represented.
