## Supplementary Figure 8 for "Unlocking the Mycorrhizal Nitrogen Pathway Puzzle: Metabolic Modelling and multi-omics unveil Pyrimidines’ Role in Maize Nutrition via Arbuscular Mycorrhizal Fungi Amidst Nitrogen Scarcity"

### Slide 1
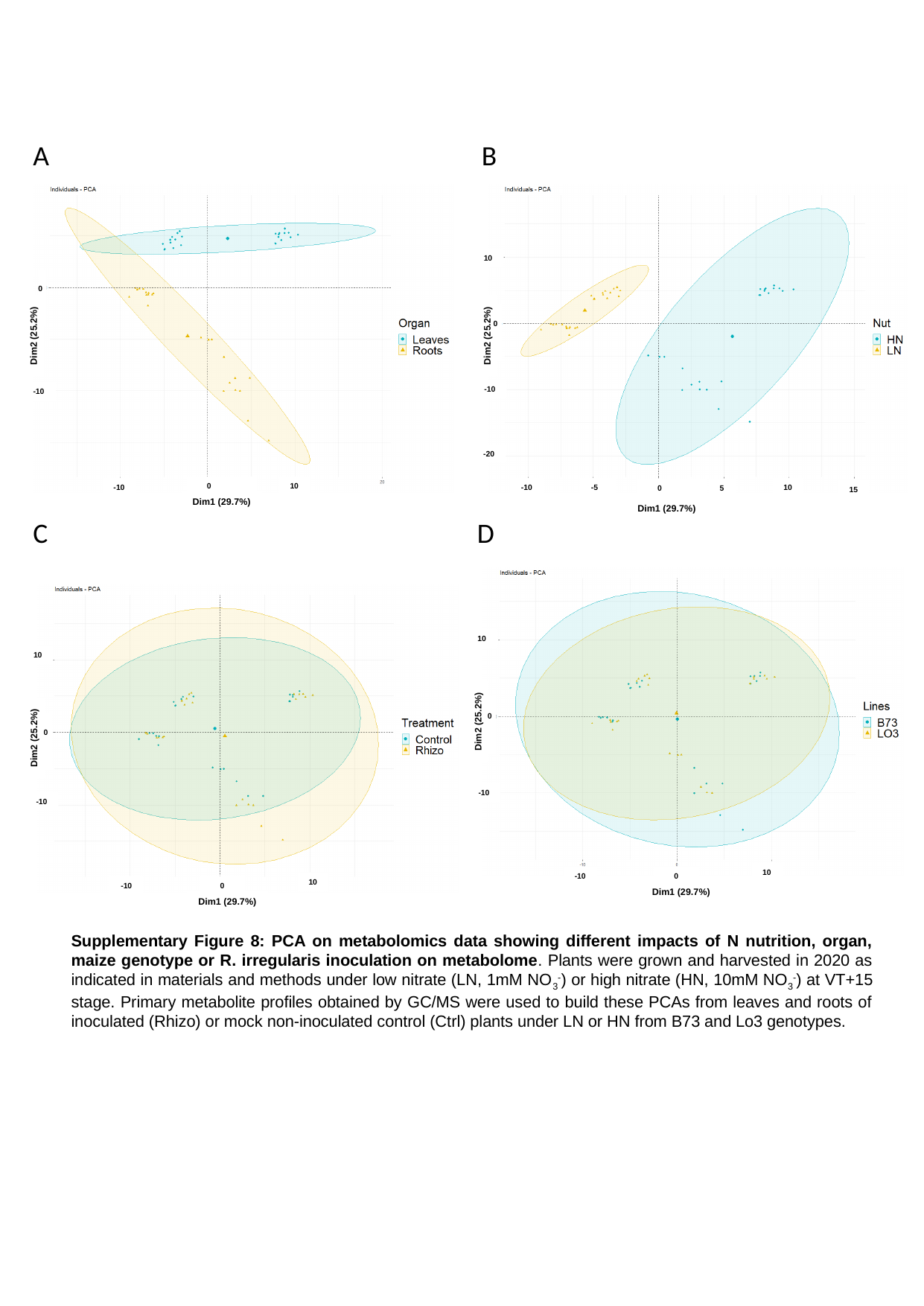

A
B
 0
Dim2 (25.2%)
-10
 0
10
-10
Dim1 (29.7%)
10
 0
Dim2 (25.2%)
-10
-20
-10
-5
10
 0
5
15
Dim1 (29.7%)
D
C
10
 0
Dim2 (25.2%)
-10
10
 0
-10
Dim1 (29.7%)
10
 0
Dim2 (25.2%)
-10
10
 0
-10
Dim1 (29.7%)
Supplementary Figure 8: PCA on metabolomics data showing different impacts of N nutrition, organ, maize genotype or R. irregularis inoculation on metabolome. Plants were grown and harvested in 2020 as indicated in materials and methods under low nitrate (LN, 1mM NO_3_^-^) or high nitrate (HN, 10mM NO_3_^-^) at VT+15 stage. Primary metabolite profiles obtained by GC/MS were used to build these PCAs from leaves and roots of inoculated (Rhizo) or mock non-inoculated control (Ctrl) plants under LN or HN from B73 and Lo3 genotypes.
