## Supplementary Figure 12 for "Unlocking the Mycorrhizal Nitrogen Pathway Puzzle: Metabolic Modelling and multi-omics unveil Pyrimidines’ Role in Maize Nutrition via Arbuscular Mycorrhizal Fungi Amidst Nitrogen Scarcity"

### Slide 1
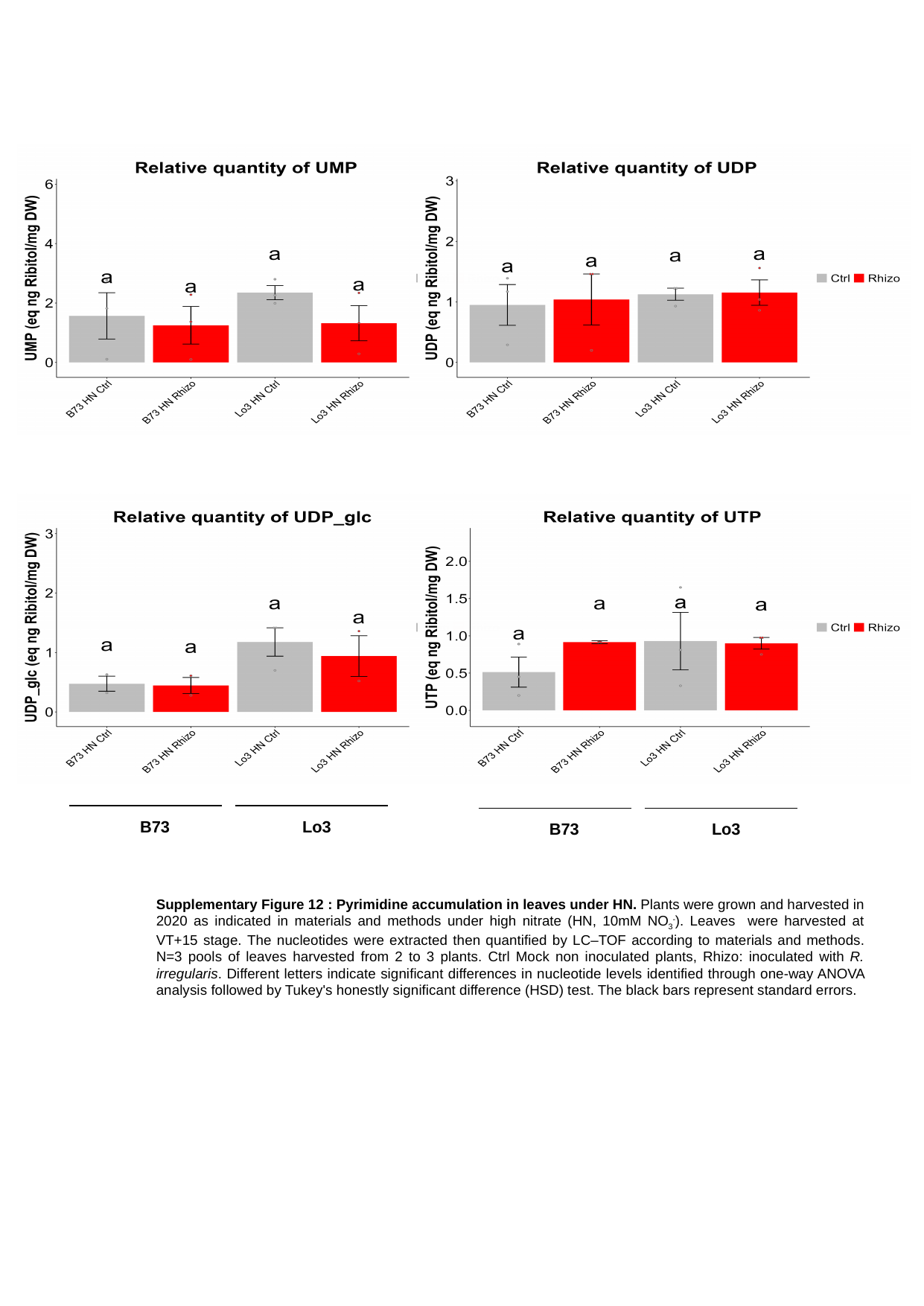

Lo3
B73
Lo3
B73
Supplementary Figure 12 : Pyrimidine accumulation in leaves under HN. Plants were grown and harvested in 2020 as indicated in materials and methods under high nitrate (HN, 10mM NO_3_^-^). Leaves were harvested at VT+15 stage. The nucleotides were extracted then quantified by LC–TOF according to materials and methods. N=3 pools of leaves harvested from 2 to 3 plants. Ctrl Mock non inoculated plants, Rhizo: inoculated with R. irregularis. Different letters indicate significant differences in nucleotide levels identified through one-way ANOVA analysis followed by Tukey's honestly significant difference (HSD) test. The black bars represent standard errors.
