## Supplementary Figure 9 for "Unlocking the Mycorrhizal Nitrogen Pathway Puzzle: Metabolic Modelling and multi-omics unveil Pyrimidines’ Role in Maize Nutrition via Arbuscular Mycorrhizal Fungi Amidst Nitrogen Scarcity"

### Slide 1
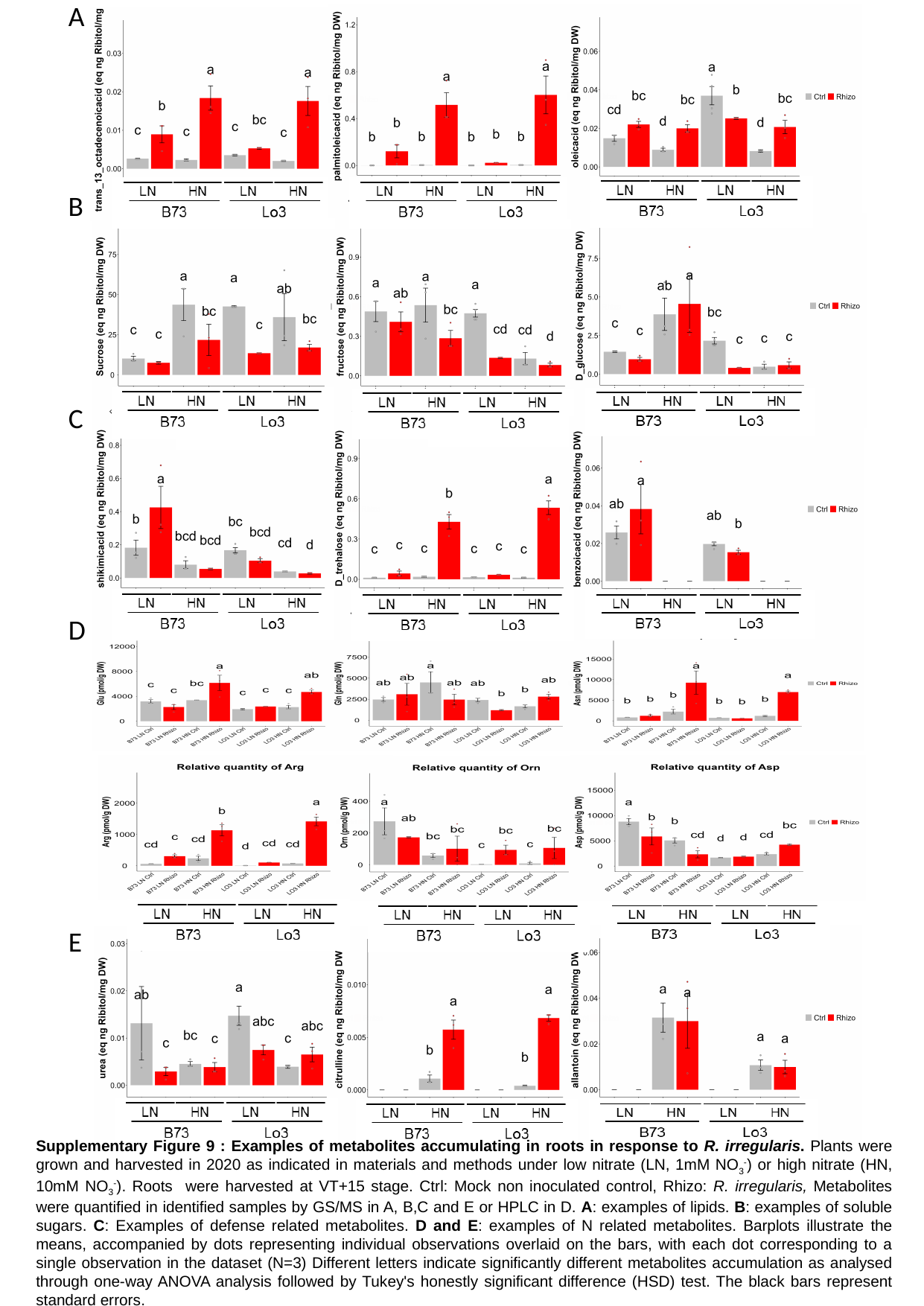

A
B
C
D
E
Supplementary Figure 9 : Examples of metabolites accumulating in roots in response to R. irregularis. Plants were grown and harvested in 2020 as indicated in materials and methods under low nitrate (LN, 1mM NO_3_^-^) or high nitrate (HN, 10mM NO_3_^-^). Roots were harvested at VT+15 stage. Ctrl: Mock non inoculated control, Rhizo: R. irregularis, Metabolites were quantified in identified samples by GS/MS in A, B,C and E or HPLC in D. A: examples of lipids. B: examples of soluble sugars. C: Examples of defense related metabolites. D and E: examples of N related metabolites. Barplots illustrate the means, accompanied by dots representing individual observations overlaid on the bars, with each dot corresponding to a single observation in the dataset (N=3) Different letters indicate significantly different metabolites accumulation as analysed through one-way ANOVA analysis followed by Tukey's honestly significant difference (HSD) test. The black bars represent standard errors.
