## Supplementary Figure 10 for "Unlocking the Mycorrhizal Nitrogen Pathway Puzzle: Metabolic Modelling and multi-omics unveil Pyrimidines’ Role in Maize Nutrition via Arbuscular Mycorrhizal Fungi Amidst Nitrogen Scarcity"

### Slide 1
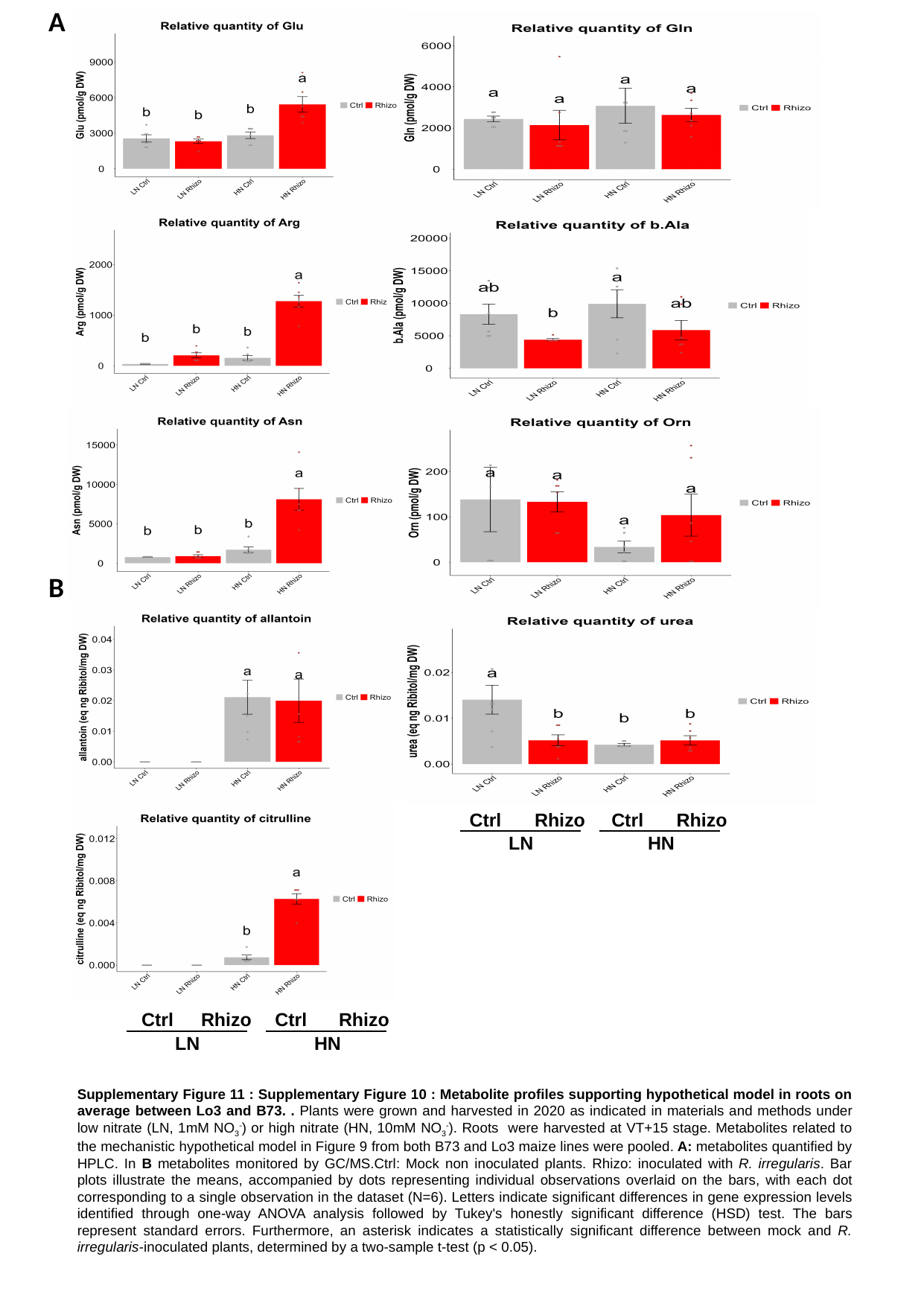

A
B
Ctrl
Rhizo
LN
Ctrl
Rhizo
HN
Ctrl
Rhizo
LN
Ctrl
Rhizo
HN
Supplementary Figure 11 : Supplementary Figure 10 : Metabolite profiles supporting hypothetical model in roots on average between Lo3 and B73. . Plants were grown and harvested in 2020 as indicated in materials and methods under low nitrate (LN, 1mM NO_3_^-^) or high nitrate (HN, 10mM NO_3_^-^). Roots were harvested at VT+15 stage. Metabolites related to the mechanistic hypothetical model in Figure 9 from both B73 and Lo3 maize lines were pooled. A: metabolites quantified by HPLC. In B metabolites monitored by GC/MS.Ctrl: Mock non inoculated plants. Rhizo: inoculated with R. irregularis. Bar plots illustrate the means, accompanied by dots representing individual observations overlaid on the bars, with each dot corresponding to a single observation in the dataset (N=6). Letters indicate significant differences in gene expression levels identified through one-way ANOVA analysis followed by Tukey's honestly significant difference (HSD) test. The bars represent standard errors. Furthermore, an asterisk indicates a statistically significant difference between mock and R. irregularis-inoculated plants, determined by a two-sample t-test (p < 0.05).
